# Pressure Points: Endothelial Responses to Shear Stress and Pressure in Health and Pulmonary Arterial Hypertension

**DOI:** 10.1101/2025.07.28.666693

**Authors:** Kellen Hirsch, Christian J. Mandrycky, Isabella Kwan, Hongyang Pi, William A. Altemeier, Tim Lahm, Peter J. Leary, Sina A. Gharib, Ying Zheng, Samuel G. Rayner

## Abstract

**Background:** Hemodynamic forces exert a profound influence on endothelial signaling and, when abnormal, contribute centrally to human vascular disease. Pulmonary arterial hypertension (PAH) is characterized by both hemodynamic derangement and pulmonary arterial endothelial cell (PAEC) dysfunction. Despite importance in disease initiation and progression, the combined effects of shear and pressure forces on PAEC biology remain incompletely understood, particularly in the context of PAH.

**Methods:** PAECs obtained at explant from controls and patients with idiopathic PAH or congenital heart disease-associated PAH (CHD-PAH) were cultured in a custom resistor-coupled microfluidic platform and exposed to static, low (3 dyne/cm^²^), or high (20 dyne/cm^²^) shear stress under either low or elevated (60 mmHg) pressure. After 24 hours, we assessed cellular morphology and performed transcriptomic analysis via bulk RNA sequencing, incorporating analyses of PAH subtype and donor sex.

**Results:** Morphologically, PAECs (n=18 donors) aligned with flow under high, but not low, shear, and alignment was not significantly altered by disease state or pressure. As expected, shear stress fundamentally reorganized the PAEC transcriptome. The “dose-response” to increasing shear differed across biological pathways in six statistically significant patterns. Increasing shear led to divergence in transcription between control and PAH cells, particularly in pathways involved in immune activation, stress signaling, and vascular remodeling, with subtype differences also observed. Pressure had modest effects on transcription, with CHD-PAH PAECs notably displaying pressure-induced stress and inflammatory signaling. We identified sexual dimorphism in the endothelial shear response, including that male cells under shear enriched for proliferative and angiogenic pathways and female cells for fatty acid metabolism and stress responses.

**Conclusions:** We provide a systems-level overview of how shear and pressure shape PAEC transcription, revealing divergent responses across disease state, PAH subtype, and donor sex. These findings highlight the need for further investigation into mechanosensitive pathways in PAH as potential novel therapeutic targets.

## Introduction

The endothelium, uniquely positioned at the interface of circulating blood and organ parenchyma, is exquisitely responsive to the hemodynamic forces imparted by blood flow. Through mechanotransduction, these forces shape fundamental endothelial programs such as alignment, proliferation, immune trafficking, and barrier integrity.^1–4^ Among them, shear stress, the tangential force exerted by flowing blood, has been the most extensively studied in endothelial biology. Within the physiologic “set point” of a given vascular bed, laminar shear stress promotes endothelial quiescence and vascular homeostasis.^3,5^ However, shear levels on either side of this physiologic range provoke endothelial activation and vascular remodeling in an attempt to restore equilibrium.^3^ Pressure induces both circumferential (hoop) and axial stress and, while pressure has been less well studied than shear stress, it has been shown to promote cellular proliferation and barrier disruption and to modulate shear-induced signaling and cellular alignment.^4,6,7^ Abnormal shear and pressure contribute to endothelial dysfunction in numerous vascular diseases including atherosclerosis, aneurysm, and pulmonary hypertension.

Pulmonary arterial hypertension (PAH) is a distinct form of pulmonary hypertension characterized by intrinsic pulmonary vasculopathy, where the effects of hemodynamic forces on pulmonary arterial endothelial cells (PAECs) are especially relevant. PAH is a progressive and currently incurable disease where increasing vascular resistance ultimately leads to intractable right heart failure.^8^ PAEC dysfunction is believed to drive key pathobiological features of PAH including inflammation, vasoconstriction, thrombosis, and intimal and medial hypertrophy.^9–12^ This disease arises via a complex interplay of patient and environmental factors, likely requiring multiple “hits.”^13^ One such environmental factor is abnormal pulmonary blood flow, which contributes etiologically in cases such as congenital heart disease-associated PAH (CHD-PAH),^14^ and potentially contributes to progression across all forms of PAH once abnormal flow patterns have developed.^11,12,15–20^ PAH is associated with a bimodal disruption of shear forces, with pathologically low shear in central, dilated arteries and distal to obstructed vasculature, and markedly elevated levels within narrow, remodeled arterioles.^21–28^ Both extremes are known to promote endothelial dysfunction.^1,11,12,16,17,29–32^ Increased vascular pressure, a requisite finding in PAH, may independently contribute to vascular pathology.^4,33^

Despite well-established clinical and basic science links between hemodynamic forces, endothelial dysfunction, and PAH, critical gaps in our understanding remain. At a fundamental level, how PAECs respond to the entire range of relevant shear stresses remains poorly defined. It is also unclear where and how the transition from physiologic to pathologic shear occurs in the pulmonary circulation. Similarly, the contribution of pressure to vascular dysfunction has been minimally studied, and the combined effects of shear stress and pressure even less so. In vivo, cells are exposed to these forces in combination, yet most studies have examined them in isolation. From a more translational standpoint, detailed transcriptomic studies exploring how shear or pressure affect PAECs in PAH at a systems level are lacking. Furthermore, it is not known if PAECs from patients with PAH respond differently to hemodynamic forces compared to controls, or how disease subtypes and patient characteristics, including biologic sex, influence these responses.

To address these gaps, we used a previously described resistor-coupled microfluidic device to evaluate PAEC responses to defined hemodynamic conditions.^4^ This platform enables a novel approach to studying the independent effects of shear and pressure on vascular cells: by adjusting the length (and thereby hydraulic resistance) of a microfluidic resistor attached to a culture channel, the pressure within the culture channel can be precisely set for a given flow rate and shear stress. In this study, we combined “low” (3 dyn/cm2) or “high” (20 dyn/cm2) shear stress with either low or high (60 mmHg) pressure, with static culture as a comparator. We first evaluated general patterns in PAEC responses to increasing shear stress at a systems level. Next, we examined how disease state and patient characteristics influence endothelial responses to hemodynamic forces using cells from patients with idiopathic PAH (IPAH), CHD-PAH, and controls. The overall goal of this effort is to provide molecular insights into how hemodynamic disturbances provoke or perpetuate PAH, in hopes of informing future development of targeted therapies to restore vascular homeostasis.

## Methods

### Design and fabrication of the resistor-coupled microfluidic

To independently modulate the pressure and shear experienced by cultured endothelial cells, we utilized a resistor-coupled microfluidic platform, as previously described.^4^ In this system, polydimethylsiloxane (PDMS) culture channels are bonded to glass slides that are then coated with polydopamine followed by fibronectin, to optimize cellular adhesion. A microfluidic pump is connected to the inlet of the culture channels via tubing, and the outlet connected to a microfluidic resistor — an empty narrow PDMS channel. By varying the length of the microfluidic resistor and the flow rate of the microfluidic pump, pressure and shear can be precisely specified (**Figure 1a**). Details on this platform, its operation, and hemodynamic calculations are provided in *Supplementary Methods*.

**Figure 1:**
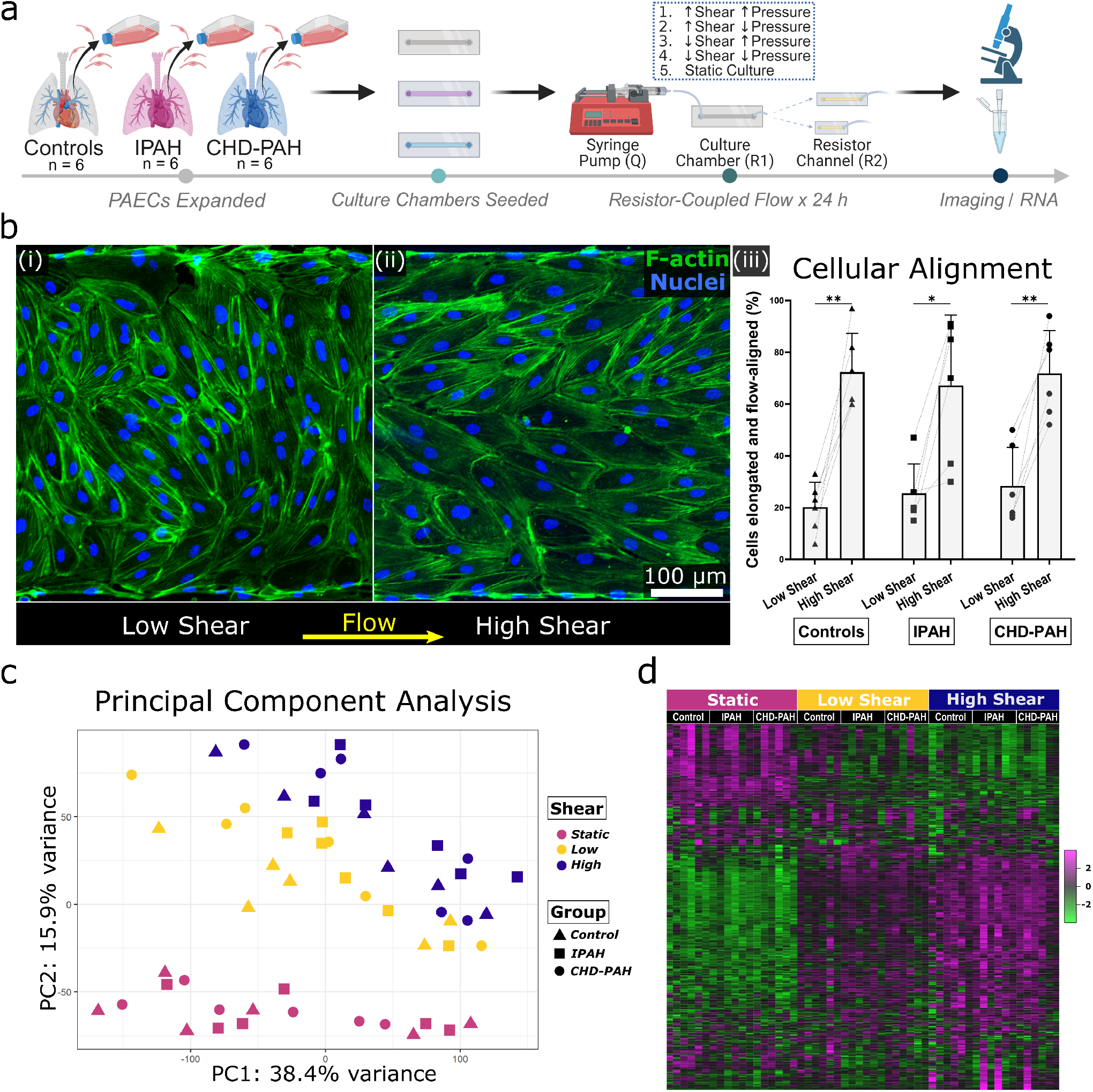
Experimental design and global morphologic and transcriptional responses to shear stress. *(a)* Timeline of experiments. Pulmonary arterial endothelial cells (PAECs) from control subjects and individuals with idiopathic pulmonary arterial hypertension (IPAH) or congenital heart disease-associated PAH (CHD-PAH) were seeded into microfluidic culture chambers, grown to confluence under creeping flow, and exposed to defined combinations of pressure and shear stress for 24 hours. Pressure and shear stress were set by varying the length of a resistor channel and adjusting the flow rate of perfused culture media. *(b)* Representative confocal images of F-actin staining in culture channels exposed to low (i) versus high (ii) shear stress for 24 hours, demonstrating flow-alignment with shear. The application of shear led to elongation and alignment within 0-30 degrees of the direction of flow (iii), without differences seen across disease states. *(c)* Principal component analysis showing transcriptional variance for all subjects under low pressure and varying shear stress. *(d)* Heatmap of transcriptional data under low pressure, organized by subject type and shear condition. ^*^ = P < 0.05, ^**^ = P<0.01.

### Endothelial cell culture

PAECs were obtained from the Pulmonary Hypertension Breakthrough Initiative (PHBI), a national consortium funded by the National Institutes of Health and the Cardiovascular Medical Research and Education Fund.^34^ PAECs from the explanted lungs of patients with idiopathic PAH (IPAH) or CHD-PAH were obtained, as well as PAECs from organ donors without PAH, whose lungs were declined for transplant (herein referred to as “controls”). PAECs were primarily obtained from small arteries <1 mm in diameter and donors from IPAH and control groups were intentionally selected to allow comparison of sex as a biological variable, with three male and three female donors in each group. Cells were received in culture from the PHBI, expanded in endothelial media (EGM2-MV, Lonza), and frozen. Cells were used between passage 5-7 throughout.

[utab]

### Perfusion of culture channels

Following cellular seeding into coated devices and overnight culture under creeping flow, channels were perfused with culture media, with 3.5% dextran added to media to simulate the viscosity of blood (∼3.5 cP). Media was degassed under vacuum for at least 30 minutes prior to perfusion. A multi-syringe infusion pump (KD Scientific) was used to deliver desired flow rates (**Table 1**) to each culture channel. A flow rate of 5.38 μl/min was calculated to provide 3 dyne/cm^2^ and flow of 35.84 μl/min to provide 20 dyne/cm^2^. Low pressure conditions did not incorporate a resistor. To generate high-pressure conditions a resistor was connected to the outlet of the culture channel and, as flow rate impacts pressure, resistors of different lengths were used for each shear-stress condition. A resistor length of 9 mm was used for the 20 dyne/cm^2^ condition and one of 62 mm for the 3 dyne/cm^2^ condition, both calculated to provide a mean pressure of 60 mmHg in the culture channel. Flow was continued for 24 hours; for the higher flow rate one syringe refill was performed 12 hours into perfusion, requiring brief interruption of flow. For the static condition, glass-bottom 24-well plates (Cellvis) were coated with 0.1% polydopamine and fibronectin as above, and cells cultured to confluence in static media. Culture time in 24-well plates was 36-48 hours, performed simultaneously with the above flow-directed channel seeding/culture/flow exposure.

**Table 1.**
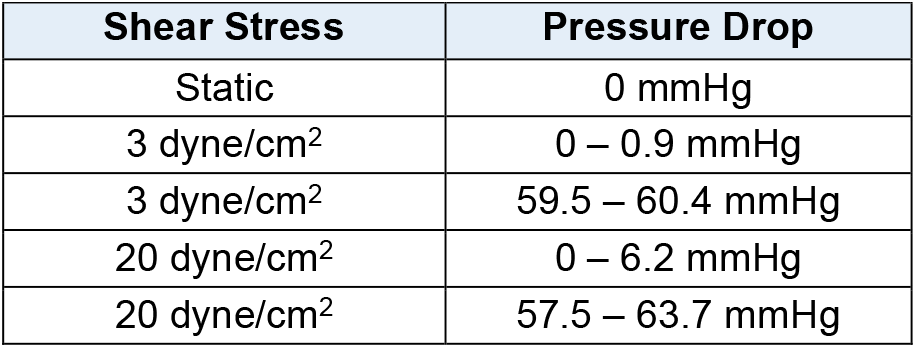
Culture Channel Parameters.

### Imaging and morphologic analysis

Brightfield images were obtained after perfusion to quantify alignment using Fiji/ImageJ.^35,36^ The percentage of cells that were both elongated and flow-aligned was compared across conditions using ANOVA with post-hoc testing. Immunofluorescent staining was done to verify endothelial identity and evaluate alignment on a subset of perfused chambers not used for RNA collection. Full protocols and details on antibodies used are provided in the *Supplementary Methods*.

### RNA isolation and sequencing

After brightfield imaging, cells were lysed in 350 μL RLT with 1% β-mercaptoethanol (Qiagen) and RNA extracted with DNase treatment (RNeasy Micro kit, Qiagen). RNA was submitted to the Fred Hutchinson Genomics Core for quality control, library preparation, and sequencing. RNA quality was confirmed via Agilent Tapestation. RNA sequencing was performed in two batches, both run in the same manner. Libraries were prepared using SMART-Seq v4 Ultra Low Input (Takara) and Nextera XT DNA Library Preparation (Illumina) kits, and sequenced on an Illumina NextSeq2000 (paired end, 50 bp). Image analysis and base calling was performed using Illumina’s Real Time Analysis v3.7.17 software, followed by demultiplexing of indexed reads and generation of FASTQ files using bcl2fastq Conversion Software v2.20 (Illumina). Alignment was performed against the GRCh38 reference genome using STAR v-2.7.1a in the two-pass alignment mode. Subread featureCounts v-1.6.0 was used to perform gene expression quantification and generate counts.

### Experimental Design Considerations

The effect of shear stress (static, low, and high) was tested across all donors. Due to feasibility constraints, the effect of high vs low pressure was assessed in a subset of donors (three each from control, CHD-PAH, and IPAH). Thus, experiments were conducted in two main batches: one batch (n = 9) exposed to all five hemodynamic conditions (shear and pressure), and another batch (n = 9) exposed only to shear conditions. RNA from each batch was purified and sequenced separately. Each round of syringe perfusion contained even numbers of donors from each condition (control, IPAH, CHD-PAH) to minimize batch effects, and RNA purification was done in one run on one single kit for each batch. For visualization of results, data were illustrated with GraphPad Prism 9 (version 9.5.1) and R package ggplot2 (version 3.4.4), with other software used as specified.

### RNA sequencing analysis

Following counts generation, filtering and batch correction were performed. Next, standard bioinformatics tools were used to perform principal-component analysis, heatmap generation, differential gene expression analysis, gene set enrichment analysis, overrepresentation analysis, and enrichment map generation. Short Time Series Expression Miner (STEM) Analysis was then performed to evaluate how gene expression changes with respect to gradual increasing shear (from static, to low, to high shear).^37^ Detailed methods for RNA sequencing analysis are provided in the *Supplementary Methods*.

## Results

### Donor and cellular characteristics

PAECs from 18 donors were obtained (**Table 2**). The mean age was 36.4 years and those with CHD-PAH tended to be older. There was a female predominance (61%), driven by the CHD-PAH group which was 83% female. Donors were predominantly White (94%) and 22% reported Hispanic/Latino ethnicity. Two patients had potential genetic causes underlying their PAH, both in the IPAH arm. For patients with CHD-PAH, atrial septal defect was the most common form of CHD (50%) followed by ventricular septal defect (33%). PAECs from donors displayed expected morphology and staining for endothelial markers von Willebrand factor and VE-Cadherin (**Figure S1**).

**Table 2.** Donor Characteristics.

| **Table 2: Donor Characteristics** |  |  |  |  |
| --- | --- | --- | --- | --- |
|  | **All Donors (n=18)** | **Controls (n=6)** | **IPAH (n=6)** | **CHD-PAH (n=6)** |
| **Age in Years** | 36.4 (13.5) | 41.7 (15.2) | 36.8 (14.8) | 47.8 (8.8) |
| **Female Sex** | 61% (11) | 50% (3) | 50% (3) | 83% (5) |
| **Race White** | 94% (17) | 83% (5) | 100% (6) | 100% (6) |
| **Race Unknown** | 6% (1) | 17% (1) | 0% (0) | 0% (0) |
| **Hispanic/Latino Ethnicity** | 22% (4) | 17% (1) | 17% (1) | 33% (2) |
| **Known Driver Mutation\*** | 11% (2) | 0% (0) | 33% (2) | 0% (0) |
| **Atrial Septal Defect** | 17% (3) | 0% (0) | 0% (0) | 50% (3) |
| **Ventricular Septal Defect** | 11% (2) | 0% (0) | 0% (0) | 33% (2) |
| **Other Congenital Systemic-Pulmonary Shunt** | 6% (1) | 0% (0) | 0% (0) | 17% (1) |

Data are presented as mean (standard deviation) or % (n). IPAH: idiopathic PAH; CHD-PAH: congenital heart disease-associated PAH. \*One patient had a known mutation in BMPR2 and one in EIF2AK4.

### Global morphologic and transcriptional responses to shear stress (all donors)

Following culture for 24 hours under defined hemodynamic conditions, cells were briefly imaged and RNA harvested, purified, and sequenced (**Figure 1a**). Initial analysis of alignment and gene expression was done for all donors under low pressure (**Figure S2**). Cells aligned in the direction of flow under high shear, but not low shear, conditions (**Figure 1b**). The percentage of cells elongated and flow-aligned for controls was 20.2 % under low shear and 72.3 % under high shear (P < 0.01); for IPAH this was 25.5 % under low shear and 67.2 % under high shear (P < 0.05); and for CHD-PAH values were 28.3 % under low shear and 71.8 % under high shear (P<0.01). There was no significant difference in alignment between disease states. Switching to the transcriptome, principal component analysis of the batch-corrected data suggested the greatest separation was related to shear exposure (static-low-high; **Figure 1c)**. Notably, while the transition from static to low shear visibly restructured the transcriptome, the shift from low shear to high shear appeared to amplify these changes rather than causing a fundamental transcriptional reorganization, as illustrated by a heat map of transcriptome-wide gene expression (**Figure 1d**).

We next examined the nature of these transcriptional changes across all endothelial cells, to gain broader understanding of the dynamics of endothelial responses to shear stress. Through DESeq2 analysis of all donors (n = 18) we identified 10,416 differentially expressed genes (DEGs) in the low shear vs static condition (5,001 upregulated and 5,415 downregulated), 11,916 DEGs in the high shear vs static condition (5,719 upregulated and 6,197 downregulated), and 6,962 DEGs in the high vs low shear condition (3,319 upregulated and 3,643 downregulated; **Figure 2a**). Looking specifically at a panel of canonical shear-sensitive genes across several functional domains selected a priori, our results correlated well with previously reported effects of shear on other EC populations (**Figure 2b**), with only a few exceptions, notably that we saw less HIF1A in low shear relative to static conditions.^38–41^ Gene set enrichment analysis (GSEA) revealed downregulation of pathways associated with proliferation under high shear stress versus static conditions, and upregulation of pathways associated with focal adhesion, cellular organization, hemostasis, and macroautophagy (**Figure 2c**). These functional themes and the extent of gene overlap were further visualized via network analysis using a Cytoscape Enrichment Map (**Figure 2d**). Short Time-series Expression Miner (STEM) analysis was performed to evaluate how genes changed longitudinally over increasing shear. Six statistically significant patterns of transcriptional response to increasing shear stress were observed (**Figure 2e**). Several pathways associated with replication and DNA repair exhibited a stepwise decrease in expression as shear increased from static, to low, to high. Some pathways associated with extracellular organization, however, were downregulated with application of low shear, without further decrease under high shear. Similarly, some pathways associated with angiogenesis, NOTCH, and muscular development were upregulated with low shear application and did not increase further under high shear, while other related pathways such as angiogenesis and oxidative stress response displayed a more linear response to increasing shear. Interestingly, there were no expression profiles where the direction of gene expression diverged between low shear and high shear (e.g. there were no “v-shaped” curves). Analysis of PAH and control samples separately reinforced these six expression patterns, suggesting they represent fundamental PAEC responses to shear.

**Figure 2:**
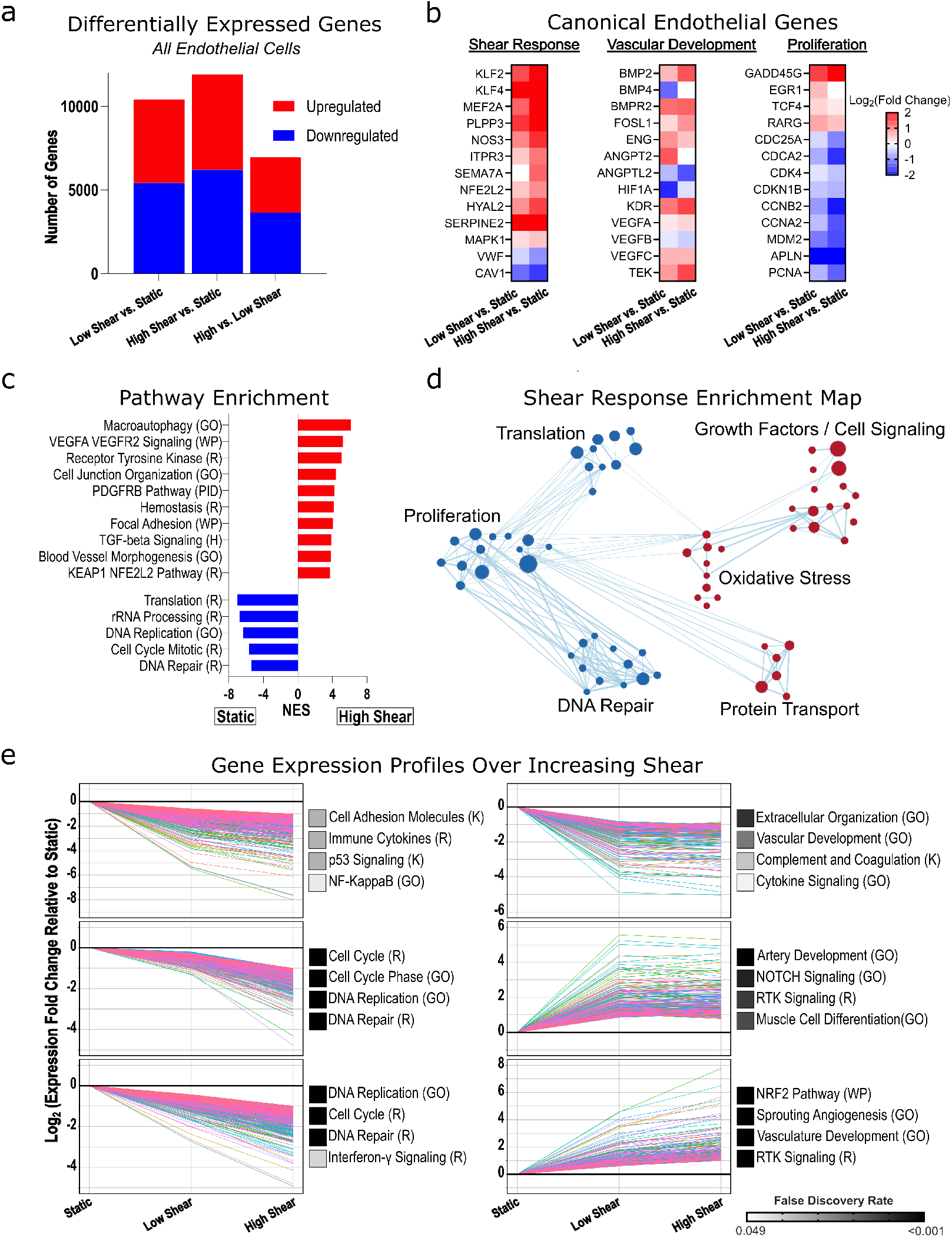
Shear stress effects on endothelial cell transcription. *(a)* Differentially expressed genes across shear conditions (n = 18, combining diseased and control subjects). *(b)* Heatmaps of log_₂_-fold change showing expected trend in select genes previously implicated in endothelial shear responses. *(c)* Gene set enrichment analysis -selected significant pathways shown, quantified by normalized enrichment score (NES). *(d)* Network of enriched gene sets in high shear (red) vs static (blue), clustered by theme. Node size reflects NES and line thickness indicates the number of overlapping genes. *(e)* Short Time-series Expression Miner (STEM) analysis identified six significant gene expression profiles. Each graph represents one expression profile, with individual lines corresponding to single genes within that profile. Profiles were functionally annotated using Gene Ontology Biological Process (GO), Kyoto Encyclopedia of Genes and Genomes (K), Reactome (R), and WikiPathways (WP). Selected pathway annotations are shown to the right, with names abbreviated for space where appropriate. Adjusted P < 0.05 used throughout.

### Endothelial responses to shear across disease states

We next examined how PAECs from PAH donors respond to increasing shear stress, compared with controls. We identified diverging gene expression between PAH (n = 12) and control (n = 6) cells with increasing shear: 11 DEGs under static conditions (7 upregulated in PAH and 4 downregulated), 71 DEGs under low shear (50 upregulated in PAH and 21 downregulated), and 180 DEGs under high shear (74 upregulated in PAH and 106 downregulated; **Figure 3a**). Under high shear, DEGs upregulated in PAH cells included those involved in vascular inflammation (MIR155, MIR155HG, ATF3, CD93, ADGRF5), proliferation and activation (PIM1 and RCAN2), and remodeling (ADAMTSL2), as well as genes associated with shear response and vascular homeostasis (e.g. GUCY1B1, CA2, and CSMD1; **Figure 3b**). In contrast, control cells showed differential upregulation of genes involved in mitochondrial function (MRPL16, TIMM22), cytoskeletal organization (ACTN2, CADM3, KCNN4), anti-inflammation/stress regulation (TNFAIP3, FST, TGFB2), and endothelial survival (FGF2, NRG1; **Figure 3b**).

**Figure 3:**
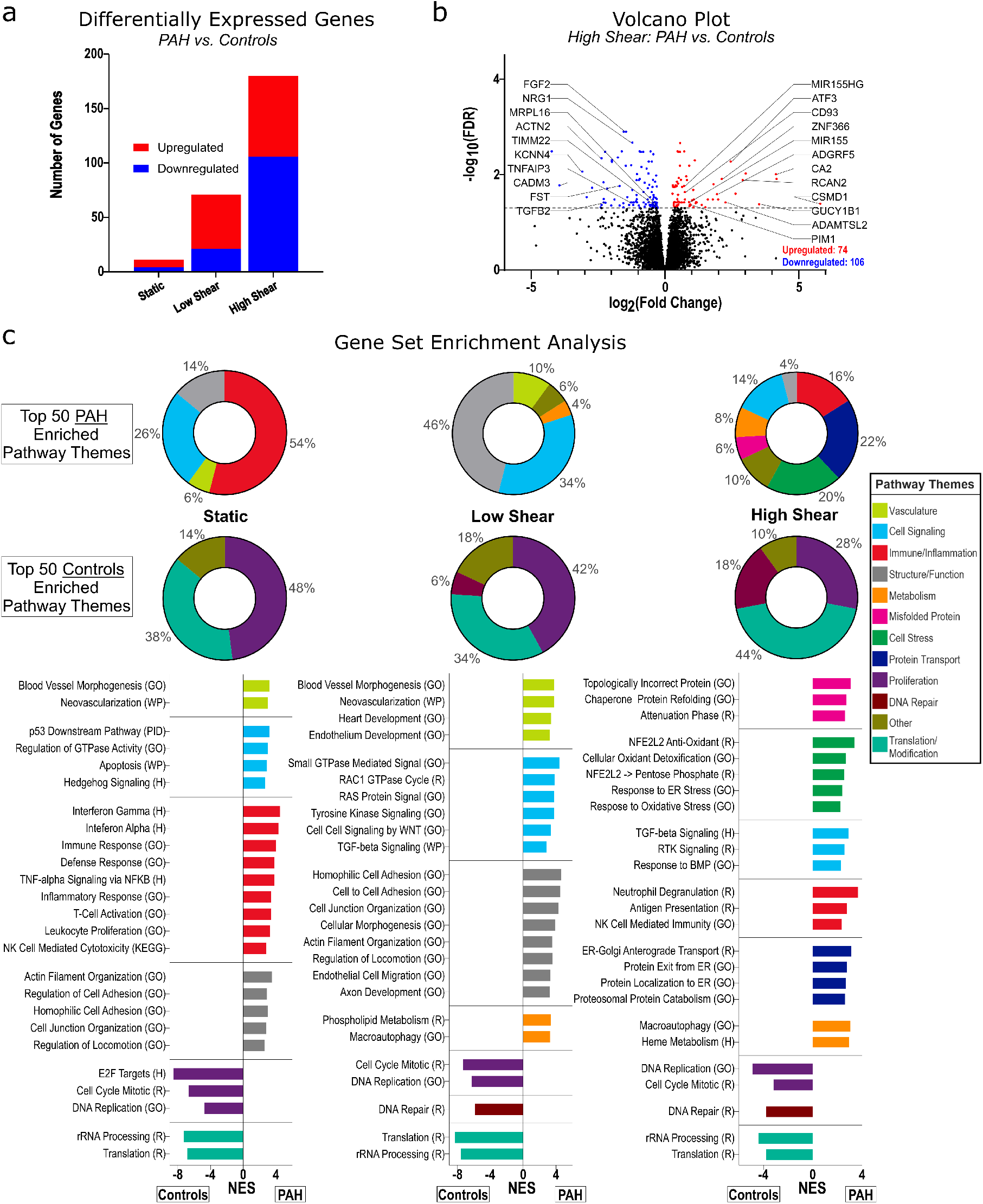
Differential responses to shear stress in PAH versus control PAECs. *(a)* Differentially expressed genes (DEGs) in PAH vs control PAECs under three shear conditions. *(b)* Volcano plot highlighting DEGs in the high shear condition; selected genes of interest labeled. *(c)* Gene set enrichment analysis comparing PAH to Control PAECs across static, low, and high shear. The top 50 enriched pathways for each condition were clustered into pre-specified themes and represented as pie charts showing the proportion of pathways in each theme. Representative enriched pathways from each theme are shown below, quantified by normalized enrichment score (NES). All analyses used FDR < 0.05 for significance and pathway annotation from Hallmark (H), KEGG (K), Reactome (R), Wikipathways (WP), and GO Biological Process (GO). Pathway names abbreviated for space where appropriate.

To identify broader transcriptional programs, we performed GSEA comparing PAH and control donors under static, low shear, and high shear conditions. The top 50 enriched gene sets for each comparison were manually categorized into 12 predefined biological themes (**Figure 3c**). Under static and low shear, PAH cells had enrichment of gene sets related to blood vessel formation, with increased inflammatory transcription under both static and high shear conditions. Notably, under high shear, PAH cells displayed increased enrichment of gene sets associated with cellular stress responses, including unfolded protein handling and autophagy. Conversely, across all shear conditions, control cells were enriched for gene sets related to translation and proliferation. To complement our global GSEA, we next performed ORA limited to DEGs, to evaluate differences in how PAH and control cells responded to high versus low shear (**Figure S3**). While similar themes emerged of decreased proliferation, cytoskeletal remodeling, vesicular trafficking, and proteostasis, PAH cells showed increased enrichment of stress signaling and immune activation, consistent with the above GSEA results comparing across conditions.

We next evaluated shear responses within specific subtypes of PAH -comparing CHD-PAH (n = 6) and IPAH (n = 6) with controls (n = 6). Comparing within each subject group, low shear vs static conditions led to 3,974 DEGs in control patients, 7,061 in IPAH cells, and 6,592 in CHD-PAH cells. High shear vs static conditions led to 6,897 DEGS in controls, 8,758 in IPAH cells, and 8,097 in CHD-PAH cells. High vs low shear produced 2,769 DEGs in controls, 3,114 in IPAH cells, and 1,504 in CHD-PAH cells (**Figure 4a**). Volcano plots highlighting key DEGs of interest in pulmonary vascular disease suggested specific shear-responsive patterns (**Figure 4b**). Across all donors, high shear was associated with downregulation of genes involved in vascular remodeling (e.g. BMP4, ANGPTL2, HIF3A) and junctional or mechanosensitive signaling (PECAM1, EPHB41L2), along with upregulation of classical endothelial shear-responsive and vascular repair genes (PDGFA/B, IL1A, KDR [encoding VEGFR2], SERPINB2, SOX17, IGF2, TGFBR3) suggestive of a homeostatic shear response. However, IPAH cells uniquely upregulated pro-angiogenic and inflammatory genes (e.g. VEGFA/JUN/PDGFD) and downregulated cell adhesion and apoptotic genes (ICAM1, CASP 6/8/10), consistent with enhanced angiogenesis and reduced cell death. CHD-PAH cells, meanwhile, uniquely downregulated EIF2AK4 and NF2, and upregulated SERPINB9, suggesting differential regulation of translational control and cell survival under shear.

**Figure 4:**
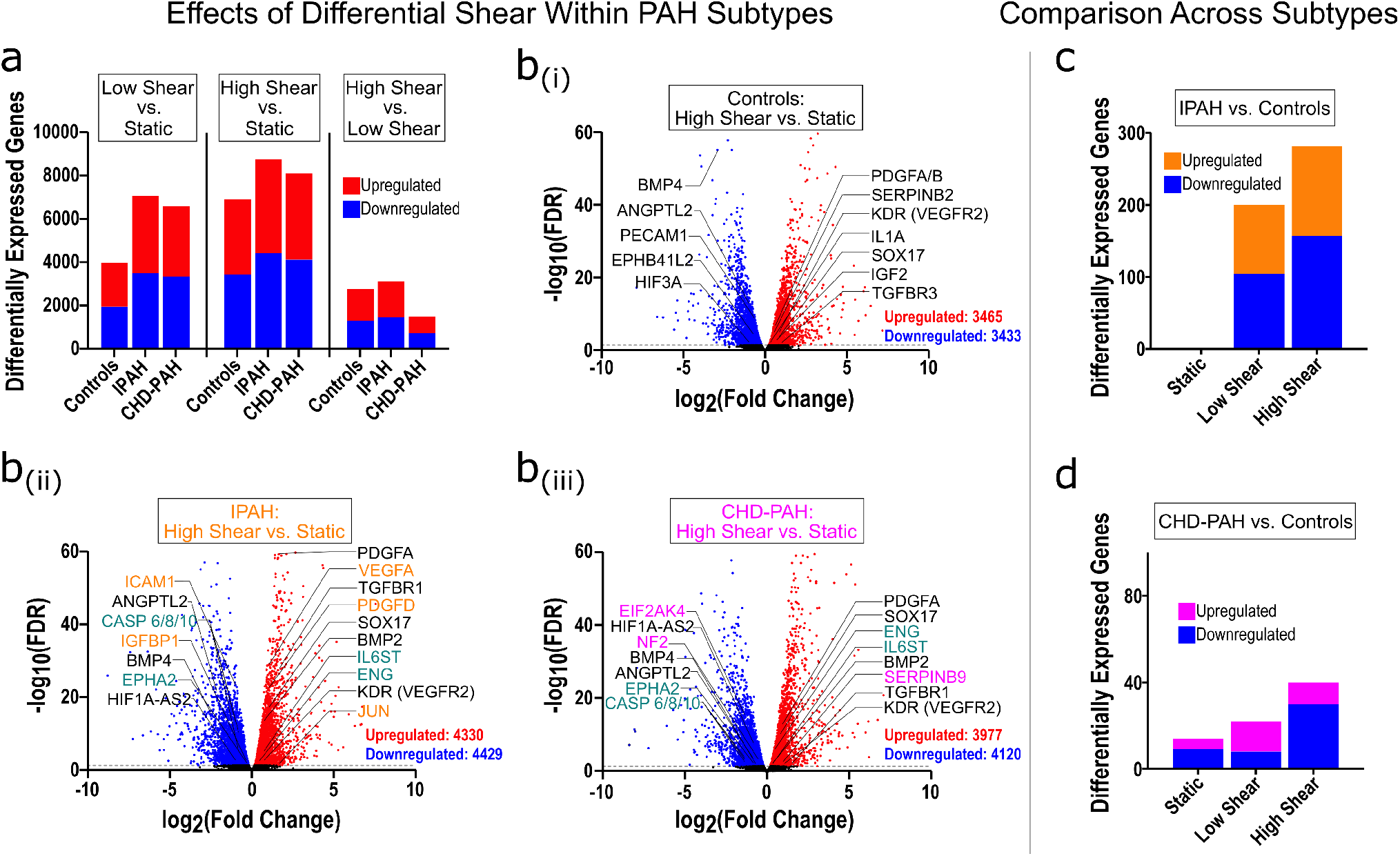
Differential transcriptional effects of shear stress across PAH subtypes. *(a)* Number of differentially expressed genes across shear conditions in PAECs from Controls (n = 6), IPAH (n = 6), and CHD-PAH (n = 6). *(b)* Volcano plots of differentially expressed genes in the high shear vs. static condition for Controls (i), IPAH (ii) and CHD-PAH (iii). Genes with significant upregulation and downregulation are colored red and blue, respectively, with select genes of interest labeled and color-coded. Genes labeled in black are differentially expressed across Controls, IPAH, and CHD-PAH, orange-labeled genes are unique to IPAH, magenta-labeled to CHD-PAH, and teal-labeled to IPAH and CHD-PAH, but not Controls. Differentially expressed genes in *(c)* IPAH vs. Controls and *(d)* CHD-PAH vs. Controls are displayed across three shear conditions. Significance is set at an adjusted p-value < 0.05 throughout.

Finally, directly comparing subtypes under each shear condition, for IPAH vs controls there was 1 DEG under static conditions, 200 DEGs under low shear, and 281 DEGs under high shear. For CHD-PAH there were 14 DEGs under static condition, 22 under low shear, and 40 under high shear (**Figure 4c+d**). Comparing CHD vs IPAH directly, there were no DEGs under static conditions or low shear conditions, and only 2 DEGs under high shear conditions (PHGHD upregulated, and TYW1B downregulated). IPAH and CHD cells showed enrichment of inflammatory and angiogenic signaling, with downregulation of proliferation and DNA-repair related pathways when compared with controls, with CHD cells under high shear displaying increased Notch4 signaling and activation of oxidative phosphorylation and metabolic signaling (**Figure S4**). Compared with IPAH, CHD cells were enriched for proliferation-related gene sets across all shear states, and under shear stress showed increased signaling related to Notch4 and RUNX1, whereas IPAH cells were enriched for extracellular matrix breakdown and signaling for muscular contraction and hypertrophy.

### The effect of high versus low pressure on shear-induced endothelial gene expression

We next sought to evaluate how the manipulation of pressure influenced cellular morphology and transcription at both low and high shear stress. A pressure of 60 mmHg was chosen as a physiologically relevant mean pressure that might be encountered in severe PAH, with the low-pressure condition reflecting the pressure imposed by flowing media in the culture channel alone with no extra resistor (ranging from 0 – 6.2 mmHg depending on flow rate and location along the culture channel). Pressure was tested in a subset of donors only (n = 3 each for control, CHD-PAH, and IPAH, respectively). One sample was a clear technical outlier within the IPAH high shear, high pressure condition and was excluded from analysis. We found that pressure did not significantly influence cellular alignment at high shear (**Figure 5a**). Principal component analysis suggested that pressure explained minimal transcriptional variance within this experiment (**Figure S5**). We also found that the application of pressure under specified states of low or high shear resulted in moderate differential gene expression (**Figure 5b)**. The greatest transcriptional change was seen for the CHD-PAH cells for high vs low pressure under low shear, where there were 29 significantly upregulated genes and 62 significantly downregulated genes. Exploring individual DEGs, we noted that pressure upregulated genes involved in cellular stress responses (e.g. GADD45B, HSP90AA1, HSPD1, and HSPB1), cytoskeletal and structural genes (e.g. KRT7 and TUBB4B), and those involved in vascular inflammation, barrier function, and angiogenesis (PROCR [EPCR] and ANGPTL4). At the same time, the application of pressure led to downregulated genes related to endothelial survival and vascular homeostasis, including MEF2C, IGFR2, and TGFBR2 (**Figure 5c)**. GSEA demonstrated that pressure generally led to enrichment in pathways for proliferation, angiogenesis, migration, and replication, with the exception of control cells under low shear where the low-pressure condition enriched for pathways related to translation and replication (**Figure S6**).

**Figure 5:**
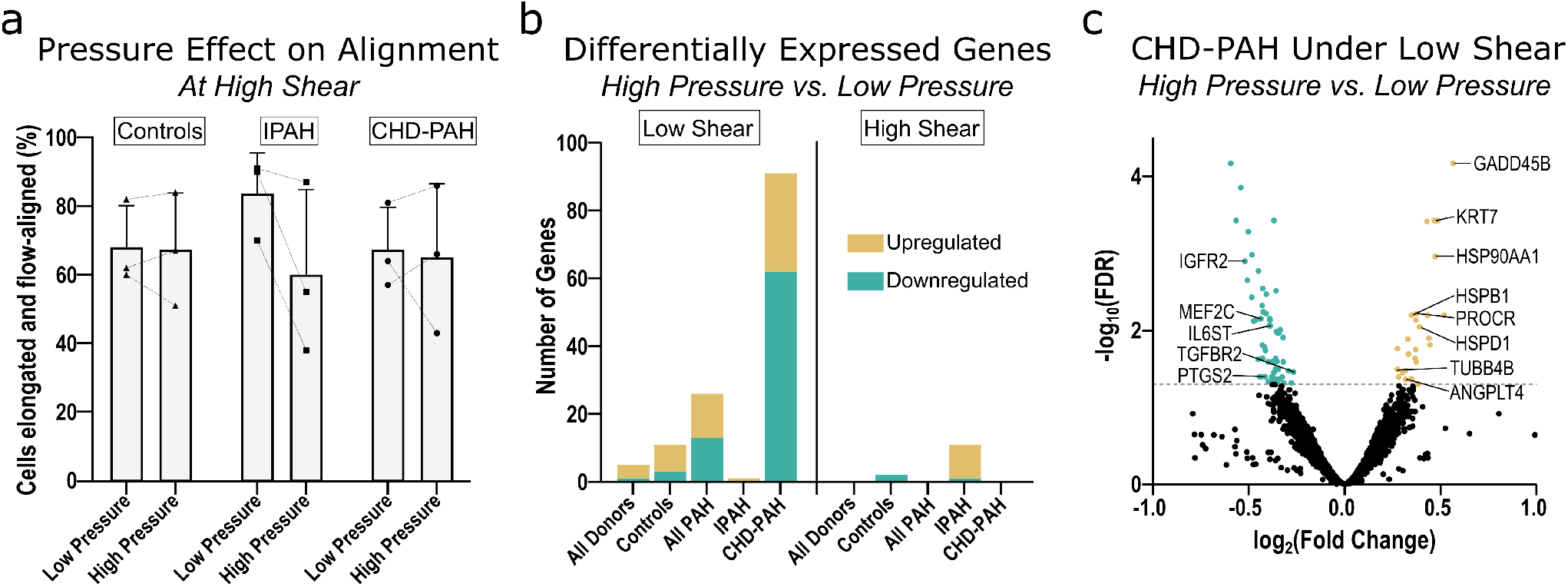
Morphologic and transcriptional responses to pressure, across shear levels and disease states: *(a)* The percentage of cells both elongated and aligned within 0-30 degrees of the direction of flow, compared across low versus high pressure conditions at high shear. Analyzed using brightfield images obtained after 24 hours of culture under shear/pressure. No comparisons were significant at P < 0.05. *(b)* Differentially expressed genes (DEGs) comparing high vs. low pressure under low shear (left) or high shear (right). Sample groups: low shear—All Donors (n = 9), Controls (n = 3), All PAH (n = 6), IPAH (n = 3), CHD-PAH (n = 3); high shear—All Donors (n = 8), Controls (n = 3), All PAH (n = 5), IPAH (n = 2), CHD-PAH (n = 3). *(c)* Volcano plot of DEGs, comparing CHD-PAH cells under high vs low pressure at low shear. Select genes of interest are labeled.

### Sex differences in the shear response

During cell sourcing from the PHBI we purposefully sex-matched the control and IPAH groups (n = 3 males and n = 3 females in each group), to be able to compare sex as a biological variable in the PAEC shear response. Comparing males to females within the IPAH and control groups, across each shear state, under static conditions there were 158 DEGs in controls and 20 in IPAH cells, under low shear conditions there were 43 DEGs in controls and 19 in IPAH cells, and under high shear conditions there were 118 DEGs in controls and 574 in IPAH cells (**Figure 6a**). Further analysis focused on static and high-shear conditions, where the greatest difference was noted in DESeq2 analysis. GSEA determined that under both static and high-shear conditions, male cells showed enrichment for pathways related to proliferation and translation, while female cells were enriched for steroid/sex-hormone signaling, and smooth muscle-related signaling (**Figure 6b**). ORA was performed on IPAH cells under high shear, and while no pathways were significantly upregulated in females, many pathways of interest were upregulated in male cells. Notably, male cells were enriched for pathways related to cellular stress responses, immune signaling, proliferation, and protein degradation, with differences also seen in vascular cytokine and sex-steroid signaling (**Figure S7)**.

**Figure 6:**
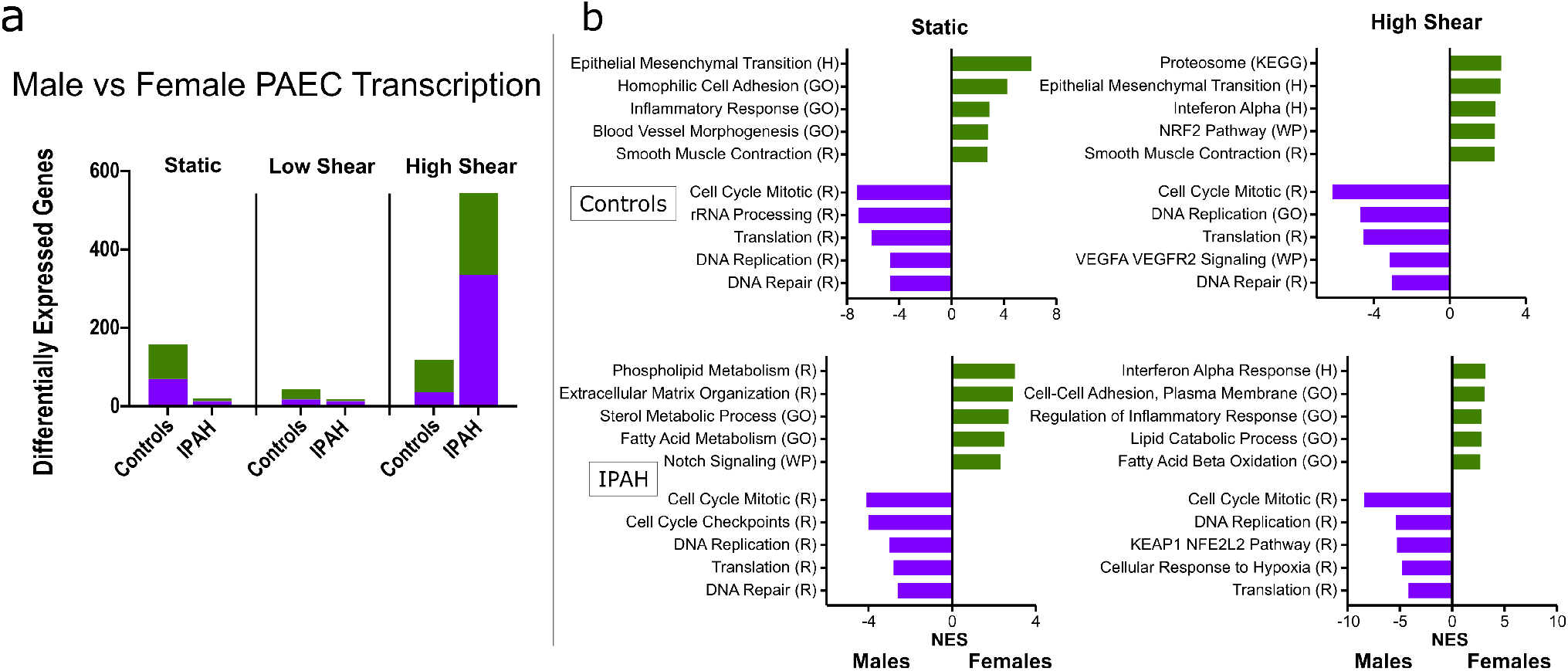
Differential transcriptional effects of shear stress on male vs female PAECs. *(a)* Differentially expressed genes (DEGs) in PAECs from females vs. males under three shear conditions. Sample groups: All Donors (n = 11 females, n = 7 males), Controls (n = 3 females, n = 3 males), and PAH (n = 3 females, n = 3 males). *(b)* Gene set enrichment analysis was used to identify significantly enriched pathways in females versus males at static (left) and high (right) shear. Significance was determined by an FDR q-value <0.05 and enrichment quantified by the normalized enrichment score (NES). Pathways are annotated with gene sets from Hallmark (H), Kyoto Encyclopedia of Genes and Genomes (KEGG), Reactome (R), Wikipathways (WP), and Gene Ontology Biological Processes (GO), with pathway names abbreviated where appropriate.

## Discussion

We conducted a systems-level analysis of PAEC responses to defined hemodynamic conditions across control and PAH donors, incorporating analyses of disease subtype and donor sex. This work provides several key insights into how shear and pressure forces shape endothelial transcriptional programs in health and disease and suggests potential future avenues for therapeutic intervention. Analyses of PAECs from PAH patients suggested distinct molecular differences across disease state and sex that are exposed by varying levels of shear stress.

First, consistent with prior studies, even low laminar shear stress led to a fundamental reorganization of the transcriptome. We found that shear’s effects on transcription were much more pronounced than those of pressure or disease state. Increasing shear from low to high levels amplified shear-induced changes in signaling, rather than provoking a fundamental shift in the direction of transcription, suggesting that shear responses exist on a continuum over the range studied here. Cluster analysis using STEM supported this concept and identified that the pulmonary endothelial response to shear is heterogeneous across biological pathways. Some pathways plateau at low shear stress, others continue to increase or decrease under higher shear, and others show an accelerating change at higher shear. While prior work, including our own preliminary evaluation of commercial PAECs at 0 and 5 dyne/cm^2^, has shown that shear stress broadly influences the endothelial transcriptome,^2,4^ to our knowledge this is the first study to systematically examine transcriptome-wide dose-dependent responses to different levels of shear stress, and to quantify heterogeneity in response at the gene and pathway level. Our use of primary organ-specific cells and biologically distinct donor replicates further increases the physiologic relevance and statistical power of our work.

Second, while PAH and control cells were transcriptionally similar at baseline, their signaling diverged significantly under hemodynamic stress. Particularly, PAH PAECs exhibited increased enrichment of cellular stress responses, immune activation, and vascular remodeling responses under high shear, suggesting impairment in the physiological adaptive response to laminar shear stress and/or an exaggerated response to hemodynamic stress – potentially contributing to PAH pathogenesis or propagation. There were differences even within subtypes of PAH, where CHD cells showed increased signaling related to Notch4 and RUNX1 and IPAH cells were enriched for extracellular matrix breakdown and signaling to smooth muscle. These differences may point to distinct pathobiology across PAH subtypes. Prior RNA sequencing-based work has also shown few differences between control and PAH cells under static conditions,^42,43^ but increased divergence in specific genes has been observed with application of chemical stimuli.^43^ We believe this to be the most comprehensive transcriptomic analysis of PAH PAEC responses to shear stress to date.

Third, while pressure had a modest effect on transcription compared to shear, CHD-PAH cells displayed increased transcriptional signals of vascular stress and inflammation under pressure. This suggests that pressure could have an amplified pathoetiologic role in certain disease subtypes, particularly in CHD where lesions are present that chronically elevate both pressure and flow. While our prior report showed impairment of flow-alignment under high pressure at 5 dyne/cm^2^, in both that work and this current study we did not see impairment of alignment at higher shear stress (20 dyne/cm^2^) – likely because this higher shear provides a strong vector for alignment that overshadows the normal pressure force.^4^

Fourth, we report preliminary evidence of sex-specific differences in endothelial signaling, both at rest and in response to shear stress. Under high shear, male PAECs showed increased proliferation and VEGF signaling, and female cells showed increased signaling for processes including fatty acid metabolism and stress responses. Previous studies had shown sexual dimorphism in flow-mediated dilation in vivo,^44^ and more recently in male versus female HUVEC morphology and YAP1 nuclear translocation with shear.^45^ Our findings are in line with clinical observations of sex differences in PAH,^46,47^ and highlight the need to consider sex differences during research into PAH with the goal of eventually developing sex-specific treatment strategies.

We chose hemodynamic conditions relevant to both healthy lungs and PAH. While pulmonary vascular pressures are routinely measured clinically, with our chosen value of 60 mmHg being consistent with advanced PAH, physiologic shear across the pulmonary vasculature remains uncertain, particularly in distal vessels. Computational fluid dynamic modeling and magnetic resonance angiography-based estimates of normal time-averaged wall shear stress in main pulmonary arteries range from ∼4 – 20 dyne/cm^2^,^21–27^ with normal values in the terminal arterioles estimated in the 20-30 dyne/cm^2^ range.^24,27^ In PAH, dichotomous disruption of shear stress is seen, with shear potentially exceeding 100 dyne/cm^2^ in stenotic arterioles and dropping below 2 dyne/cm^2^ in dilated central vessels under low cardiac output, or in hypoperfused areas distal to obstructed arterioles.^21–28^ We selected 3 dyne/cm^2^ to model pathologically low shear and 20 dyne/cm^2^ for the upper range of physiologic shear. The lack of PAEC alignment at 3 dyne/cm^2^ supports this being a pathologically low value. Our findings suggest that, rather than discrete pathologic thresholds, the effects of shear stress exist on a continuum across the range studied, with the dose-response differing for different pathways. This may break down at the extremes: prior work showed a reversal in shear-induced signaling in some cases at 100 dyne/cm,^2^ including reduced ERG and BMPR2 expression.^48^ In testing our system, shear this high led to cellular damage and delamination, precluding similar analysis. Future work exploring these extremes will be informative.

Abnormal shear stress promotes PAEC dysfunction relevant to PAH, including abnormalities in proliferation, apoptosis, nitric oxide and prostacyclin signaling, caveolin-1, endothelial-mesenchymal transition, and PPAR-γ signaling.^1,11,12,16,17,29–32^ In many instances, abnormal blood flow may be the chief stimulus leading to pulmonary vascular dysfunction, including PAH-CHD,^49^ pneumonectomy,^50^ absence of a pulmonary artery,^51^ and chronic thromboembolic pulmonary hypertension (where blood flow is redistributed to nonoccluded areas).^52^ Reduction of abnormal flow via pulmonary artery banding in animal models, or unilateral lung transplant in humans, leads to regression of PAH features.^19,20^ Although ∼10% of CHD patients develop PAH, patients with similar lesions have differing disease penetrance, suggesting that patient-specific genetic or epigenetic factors influence individual sensitivity to hemodynamic stress.^14,53,54^

Our study has several unique strengths. This includes a high number of biological replicates for an in vitro bioengineered endothelial model (n = 18), employment of precise combinations of shear and pressure, and consideration of disease subtypes and donor sex. Several important limitations also exist. First, while increased pressure leads to wall strain in vivo, this component was missing from our studies performed in a rigid culture chamber.^27^ Second, we used steady flow, rather than the pulsatile flow experienced in the pulmonary arteries in vivo. Third, the flow profile in rectangular channels is not uniform, varying from maximal shear in the center of the channel to near-zero at the walls. Fourth, while we provide a broad overview of the transcriptional landscape in response to hemodynamic forces, protein-level validation will be an important next step. Fifth, given the central importance of EC-SMC signaling in PAH pathobiology, future models incorporating SMCs will be important. Sixth, as the CHD-PAH group was predominantly female, sex is a confounder for differences observed in this subgroup. Finally, we used cells obtained from patients with end-stage PAH, which were expanded in culture prior to being used in our experiments. We unexpectedly observed less proliferative signaling in the PAH cells compared with controls, which may be related to the presence of senescent cells in this setting.

In summary, hemodynamic forces are significantly altered in PAH, may incite disease in some cases, and may lead to progression in all patients with PAH. Our findings highlight that PAEC responses to these forces are complex, varying by disease state and sex, with transcriptional thresholds and dose-responses varying across molecular pathways. The differential sensitivity to shear and pressure observed in PAH suggests that targeted modulation of mechanosensitive pathways may provide a new therapeutic approach in PAH. Future work incorporating cyclical strain, multiple cell types, and protein-level analyses will be critical next steps.

## List of non-standard abbreviations

BMPR2: Bone morphogenic protein receptor type 2
CHD-PAH: Congenital heart disease-associated PAH
IPAH: Idiopathic PAH
mPAP: Mean pulmonary artery pressure
PAH: Pulmonary arterial hypertension
PAEC: Pulmonary arterial endothelial cell
PHBI: Pulmonary Hypertension Breakthrough Initiative

## Data Availability

Raw and processed data files have been uploaded to Geo, accession number GSE303084, and will be made publicly available upon publication.

## Acknowledgments

We greatly appreciate the support of the Pulmonary Hypertension Breakthrough Initiative (PHBI), from which donor cells were sourced. The PHBI is funded by the National Institutes of Health (NIH) R24HL123767 and the Cardiovascular Medical Research and Education Fund (CMREF) UL1RR024986. Additionally, part of this work was conducted at the Washington Nanofabrication Facility / Molecular Analysis Facility, a National Nanotechnology Coordinated Infrastructure (NNCI) site at the University of Washington with partial support from the National Science Foundation via awards NNCI-2025489 and NNCI-1542101.

## Citations

1. Happé CM, Szulcek R, Voelkel NF, Bogaard HJ. Reconciling paradigms of abnormal pulmonary blood flow and quasi-malignant cellular alterations in pulmonary arterial hypertension. Vascular pharmacology. 2016;83:17–25.

2. Ajami NE, Gupta S, Maurya MR, Nguyen P, Li JY-S, Shyy JYJ, Chen Z, Chien S, Subramaniam S. Systems biology analysis of longitudinal functional response of endothelial cells to shear stress. Proceedings of the National Academy of Sciences of the United States of America. 2017;114(41):10990–10995.

3. Baeyens N, Nicoli S, Coon BG, Ross TD, Van den Dries K, Han J, Lauridsen HM, Mejean CO, Eichmann A, Thomas J-L, Humphrey JD, Schwartz MA. Vascular remodeling is governed by a VEGFR3-dependent fluid shear stress set point. eLife. 2015;4(4):1–35.

4. Mandrycky C, Ishida T, Rayner SG, Heck AM, Hadland B, Zheng Y. Under pressure: integrated endothelial cell response to hydrostatic and shear stresses. bioRxiv : the preprint server for biology. 2024.

5. Baeyens N, Bandyopadhyay C, Coon BG, Yun S, Schwartz MA. Endothelial fluid shear stress sensing in vascular health and disease. The Journal of clinical investigation. 2016;126(3):821–8.

6. Schwartz EA, Bizios R, Medow MS, Gerritsen ME. Exposure of human vascular endothelial cells to sustained hydrostatic pressure stimulates proliferation. Involvement of the alphaV integrins. Circulation research. 1999;84(3):315–22.

7. Prystopiuk V, Fels B, Simon CS, Liashkovich I, Pasrednik D, Kronlage C, Wedlich-Söldner R, Oberleithner H, Fels J. A two-phase response of endothelial cells to hydrostatic pressure. Journal of cell science. 2018;131(12).

8. Thenappan T, Ormiston ML, Ryan JJ, Archer SL. Pulmonary arterial hypertension: pathogenesis and clinical management. BMJ (Clinical research ed.). 2018;360:j5492.

9. Tuder RM, Marecki JC, Richter A, Fijalkowska I, Flores S. Pathology of pulmonary hypertension. Clinics in chest medicine. 2007;28(1):23–42, vii.

10. Budhiraja R, Tuder RM, Hassoun PM. Endothelial dysfunction in pulmonary hypertension. Circulation. 2004;109(2):159–65.

11. Humbert M, Guignabert C, Bonnet S, Dorfmüller P, Klinger JR, Nicolls MR, Olschewski AJ, Pullamsetti SS, Schermuly RT, Stenmark KR, Rabinovitch M. Pathology and pathobiology of pulmonary hypertension: state of the art and research perspectives. The European respiratory journal. 2019;53(1).

12. Ranchoux B, Harvey LD, Ayon RJ, Babicheva A, Bonnet S, Chan SY, Yuan JXJ, Perez V de J. Endothelial dysfunction in pulmonary arterial hypertension: An evolving landscape (2017 Grover Conference Series). Pulmonary Circulation. 2018;8(1).

13. Yuan JX-J, Rubin LJ. Pathogenesis of pulmonary arterial hypertension: the need for multiple hits. Circulation. 2005;111(5):534–8.

14. D’Alto M, Mahadevan VS. Pulmonary arterial hypertension associated with congenital heart disease. European respiratory review : an official journal of the European Respiratory Society. 2012;21(126):328–37.

15. Szulcek R, Happé CM, Rol N, Fontijn RD, Dickhoff C, Hartemink KJ, Grünberg K, Tu L, Timens W, Nossent GD, Paul MA, Leyen TA, Horrevoets AJ, de Man FS, Guignabert C, et al. Delayed Microvascular Shear Adaptation in Pulmonary Arterial Hypertension. Role of Platelet Endothelial Cell Adhesion Molecule-1 Cleavage. American journal of respiratory and critical care medicine. 2016;193(12):1410–20.

16. Frank PG, Lisanti MP. Role of caveolin-1 in the regulation of the vascular shear stress response. The Journal of clinical investigation. 2006;116(5):1222–5.

17. Moonen J-RAJ, Lee ES, Schmidt M, Maleszewska M, Koerts JA, Brouwer LA, van Kooten TG, van Luyn MJA, Zeebregts CJ, Krenning G, Harmsen MC. Endothelial-to-mesenchymal transition contributes to fibro-proliferative vascular disease and is modulated by fluid shear stress. Cardiovascular research. 2015;108(3):377–86.

18. Kuebler WM. Effects of Pressure and Flow on the Pulmonary Endothelium.; 2009.

19. Abe K, Shinoda M, Tanaka M, Kuwabara Y, Yoshida K, Hirooka Y, McMurtry IF, Oka M, Sunagawa K. Haemodynamic unloading reverses occlusive vascular lesions in severe pulmonary hypertension. Cardiovascular research. 2016;111(1):16–25.

20. Levy NT, Liapis H, Eisenberg PR, Botney MD, Trulock EP. Pathologic regression of primary pulmonary hypertension in left native lung following right single-lung transplantation. The Journal of heart and lung transplantation : the official publication of the International Society for Heart Transplantation. 2001;20(3):381–4.

21. Tang BT, Pickard SS, Chan FP, Tsao PS, Taylor CA, Feinstein JA. Wall shear stress is decreased in the pulmonary arteries of patients with pulmonary arterial hypertension: An image-based, computational fluid dynamics study. Pulmonary circulation. 2012;2(4):470– 6.

22. Truong U, Fonseca B, Dunning J, Burgett S, Lanning C, Ivy DD, Shandas R, Hunter K, Barker AJ. Wall shear stress measured by phase contrast cardiovascular magnetic resonance in children and adolescents with pulmonary arterial hypertension. Journal of cardiovascular magnetic resonance : official journal of the Society for Cardiovascular Magnetic Resonance. 2013;15(1):81.

23. Barker AJ, Roldán-Alzate A, Entezari P, Shah SJ, Chesler NC, Wieben O, Markl M, François CJ. Four-dimensional flow assessment of pulmonary artery flow and wall shear stress in adult pulmonary arterial hypertension: results from two institutions. Magnetic resonance in medicine. 2015;73(5):1904–13.

24. Yang W, Dong M, Rabinovitch M, Chan FP, Marsden AL, Feinstein JA. Evolution of hemodynamic forces in the pulmonary tree with progressively worsening pulmonary arterial hypertension in pediatric patients. Biomechanics and modeling in mechanobiology. 2019;18(3):779–796.

25. Zambrano BA, McLean NA, Zhao X, Tan J, Zhong L, Figueroa CA, Lee LC, Baek S. Image-based computational assessment of vascular wall mechanics and hemodynamics in pulmonary arterial hypertension patients. Journal of biomechanics. 2018;68:84–92.

26. Schäfer M, Ivy DD, Barker AJ, Kheyfets V, Shandas R, Abman SH, Hunter KS, Truong U. Characterization of CMR-derived haemodynamic data in children with pulmonary arterial hypertension. European heart journal. Cardiovascular Imaging. 2017;18(4):424–431.

27. Bartolo MA, Qureshi MU, Colebank MJ, Chesler NC, Olufsen MS. Numerical predictions of shear stress and cyclic stretch in pulmonary hypertension due to left heart failure. Biomechanics and modeling in mechanobiology. 2022;21(1):363–381.

28. Kheyfets VO, Rios L, Smith T, Schroeder T, Mueller J, Murali S, Lasorda D, Zikos A, Spotti J, Reilly JJ, Finol EA. Patient-specific computational modeling of blood flow in the pulmonary arterial circulation. Computer Methods and Programs in Biomedicine. 2015;120(2):88–101.

29. Mack JJ, Mosqueiro TS, Archer BJ, Jones WM, Sunshine H, Faas GC, Briot A, Aragón RL, Su T, Romay MC, McDonald AI, Kuo C-H, Lizama CO, Lane TF, Zovein AC, et al. NOTCH1 is a mechanosensor in adult arteries. Nature communications. 2017;8(1):1620.

30. Davies PF. Hemodynamic shear stress and the endothelium in cardiovascular pathophysiology. Nature clinical practice. Cardiovascular medicine. 2009;6(1):16–26.

31. Dabral S, Tian X, Kojonazarov B, Savai R, Ghofrani HA, Weissmann N, Florio M, Sun J, Jonigk D, Maegel L, Grimminger F, Seeger W, Savai Pullamsetti S, Schermuly RT. Notch1 signalling regulates endothelial proliferation and apoptosis in pulmonary arterial hypertension. The European respiratory journal. 2016;48(4):1137–1149.

32. Ameshima S, Golpon H, Cool CD, Chan D, Vandivier RW, Gardai SJ, Wick M, Nemenoff RA, Geraci MW, Voelkel NF. Peroxisome proliferator-activated receptor gamma (PPARgamma) expression is decreased in pulmonary hypertension and affects endothelial cell growth. Circulation research. 2003;92(10):1162–9.

33. Al-Nuaimi DA, Rütsche D, Abukar A, Hiebert P, Zanetti D, Cesarovic N, Falk V, Werner S, Mazza E, Giampietro C. Hydrostatic pressure drives sprouting angiogenesis via adherens junction remodelling and YAP signalling. Communications biology. 2024;7(1):940.

34. Stacher E, Graham BB, Hunt JM, Gandjeva A, Groshong SD, McLaughlin V V, Jessup M, Grizzle WE, Aldred MA, Cool CD, Tuder RM. Modern age pathology of pulmonary arterial hypertension. American journal of respiratory and critical care medicine. 2012;186(3):261–72.

35. Schneider CA, Rasband WS, Eliceiri KW. NIH Image to ImageJ: 25 years of image analysis. Nature methods. 2012;9(7):671–5.

36. Schindelin J, Arganda-Carreras I, Frise E, Kaynig V, Longair M, Pietzsch T, Preibisch S, Rueden C, Saalfeld S, Schmid B, Tinevez J-Y, White DJ, Hartenstein V, Eliceiri K, Tomancak P, et al. Fiji: an open-source platform for biological-image analysis. Nature methods. 2012;9(7):676–82.

37. Ernst J, Bar-Joseph Z. STEM: a tool for the analysis of short time series gene expression data. BMC bioinformatics. 2006;7:191.

38. Atkins GB, Jain MK. Role of Krüppel-like transcription factors in endothelial biology. Circulation research. 2007;100(12):1686–95.

39. Tamargo IA, Baek KI, Kim Y, Park C, Jo H. Flow-induced reprogramming of endothelial cells in atherosclerosis. Nature reviews. Cardiology. 2023;20(11):738–753.

40. Maurya MR, Gupta S, Li JYS, Ajami NE, Chen ZB, Shyy JYJ, Chien S, Subramaniam S. Longitudinal shear stress response in human endothelial cells to atheroprone and atheroprotective conditions. Proceedings of the National Academy of Sciences of the United States of America. 2021;118(4):2–9.

41. Akimoto S, Mitsumata M, Sasaguri T, Yoshida Y. Laminar shear stress inhibits vascular endothelial cell proliferation by inducing cyclin-dependent kinase inhibitor p21(Sdi1/Cip/Waf1). Circulation Research. 2000;86(2):185–190.

42. Rhodes CJ, Im H, Cao A, Hennigs JK, Wang L, Sa S, Chen PI, Nickel NP, Miyagawa K, Hopper RK, Tojais NF, Li CG, Gu M, Spiekerkoetter E, Xian Z, et al. RNA sequencing analysis detection of a novel pathway of endothelial dysfunction in pulmonary arterial hypertension. American Journal of Respiratory and Critical Care Medicine. 2015;192(3):356–366.

43. Reyes-Palomares A, Gu M, Grubert F, Berest I, Sa S, Kasowski M, Arnold C, Shuai M, Srivas R, Miao S, Li D, Snyder MP, Rabinovitch M, Zaugg JB. Remodeling of active endothelial enhancers is associated with aberrant gene-regulatory networks in pulmonary arterial hypertension. Nature Communications. 2020;11(1):1–14.

44. Tremblay JC, Stimpson T V., Pyke KE. Evidence of sex differences in the acute impact of oscillatory shear stress on endothelial function. Journal of applied physiology (Bethesda, Md. : 1985). 2019;126(2):314–321.

45. James BD, Allen JB. Sex-Specific Response to Combinations of Shear Stress and Substrate Stiffness by Endothelial Cells In Vitro. Advanced healthcare materials. 2021;10(18):e2100735.

46. Ventetuolo CE, Hess E, Austin ED, Barón AE, Klinger JR, Lahm T, Maddox TM, Plomondon ME, Thompson L, Zamanian RT, Choudhary G, Maron BA. Sex-based differences in veterans with pulmonary hypertension: Results from the veterans affairs-clinical assessment reporting and tracking database. PloS one. 2017;12(11):e0187734.

47. DesJardin JT, Kime N, Kolaitis NA, Kronmal RA, Lammi MR, Mathai SC, Ventetuolo CE, De Marco T, PHAR Investigators. Investigating the “sex paradox” in pulmonary arterial hypertension: Results from the Pulmonary Hypertension Association Registry (PHAR). The Journal of heart and lung transplantation : the official publication of the International Society for Heart Transplantation. 2024;43(6):901–910.

48. Shinohara T, Moonen J, Chun YH, Lee-Yow YC, Okamura K, Szafron JM, Kaplan J, Cao A, Wang L, Guntur D, Taylor S, Isobe S, Dong M, Yang W, Guo K, et al. High Shear Stress Reduces ERG Causing Endothelial-Mesenchymal Transition and Pulmonary Arterial Hypertension. Arteriosclerosis, thrombosis, and vascular biology. 2024;(February):1–20.

49. Dimopoulos K, Wort SJ, Gatzoulis MA. Pulmonary hypertension related to congenital heart disease: a call for action. European heart journal. 2014;35(11):691–700.

50. Potaris K, Athanasiou A, Konstantinou M, Zaglavira P, Theodoridis D, Syrigos KN. Pulmonary hypertension after pneumonectomy for lung cancer. Asian cardiovascular & thoracic annals. 2014;22(9):1072–9.

51. Ten Harkel ADJ, Blom NA, Ottenkamp J. Isolated unilateral absence of a pulmonary artery: a case report and review of the literature. Chest. 2002;122(4):1471–7.

52. Simonneau G, Torbicki A, Dorfmüller P, Kim N. The pathophysiology of chronic thromboembolic pulmonary hypertension. European respiratory review : an official journal of the European Respiratory Society. 2017;26(143):S215–S221.

53. Simonneau G, Galiè N, Rubin LJ, Langleben D, Seeger W, Domenighetti G, Gibbs S, Lebrec D, Speich R, Beghetti M, Rich S, Fishman A. Clinical classification of pulmonary hypertension. Journal of the American College of Cardiology. 2004;43(12 Suppl S):5S–12S.

54. Pascall E, Tulloh RM. Pulmonary hypertension in congenital heart disease. Future cardiology. 2018;14(4):343–353.

