## Supplementary Figures for "Pressure Points: Endothelial Responses to Shear Stress and Pressure in Health and Pulmonary Arterial Hypertension"

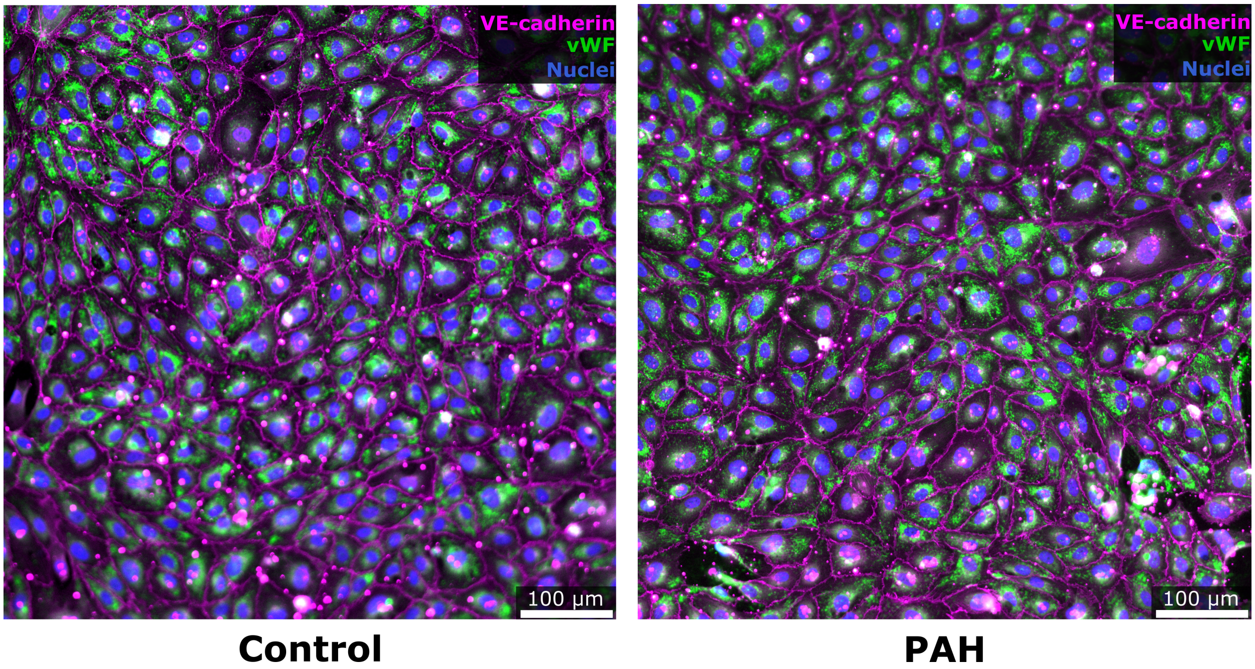


**Figure S1: Immunofluorescence of cultured subject PAECs.** Representative image of Control (left) and PAH (right) patient cells cultured to confluence and stained for nuclei (blue) and the endothelial markers VE-Cadherin (magenta) and von Willebrand factor (green).

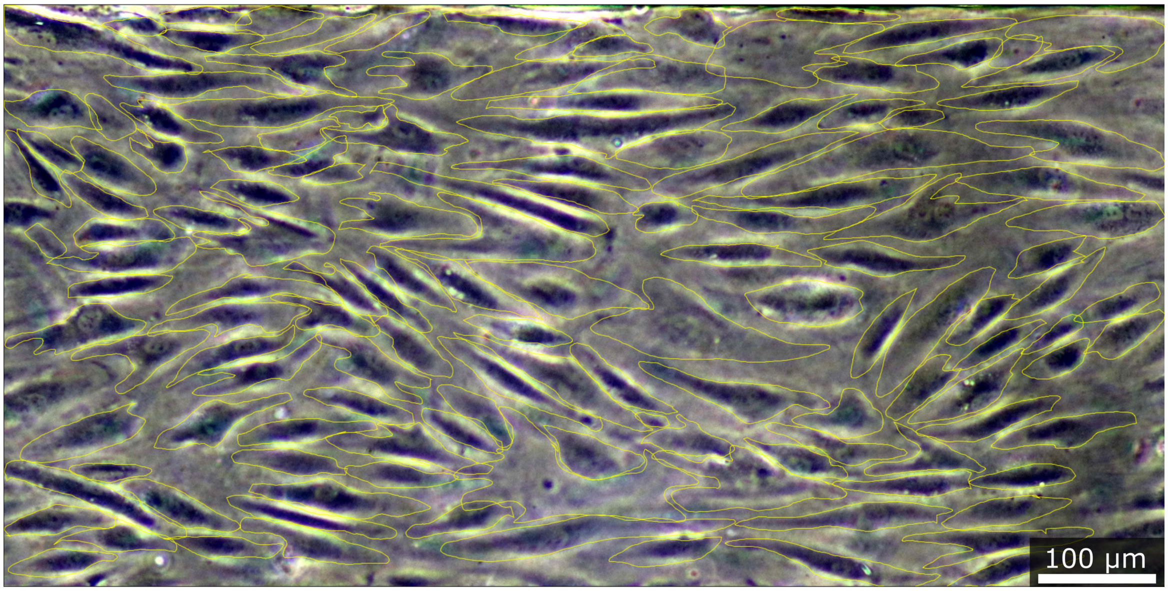


**Figure S2: PAEC alignment quantification from brightfield images.** Brightfield images were captured after 24 hours of culture under specified shear and pressure conditions. Approximate cell outlines were traced in ImageJ, and the circularity and alignment angle of each cell were measured. Cells were considered elongated and flow-aligned if their circularity was <0.7 and their alignment angle between -30° and 30° of the direction of flow.

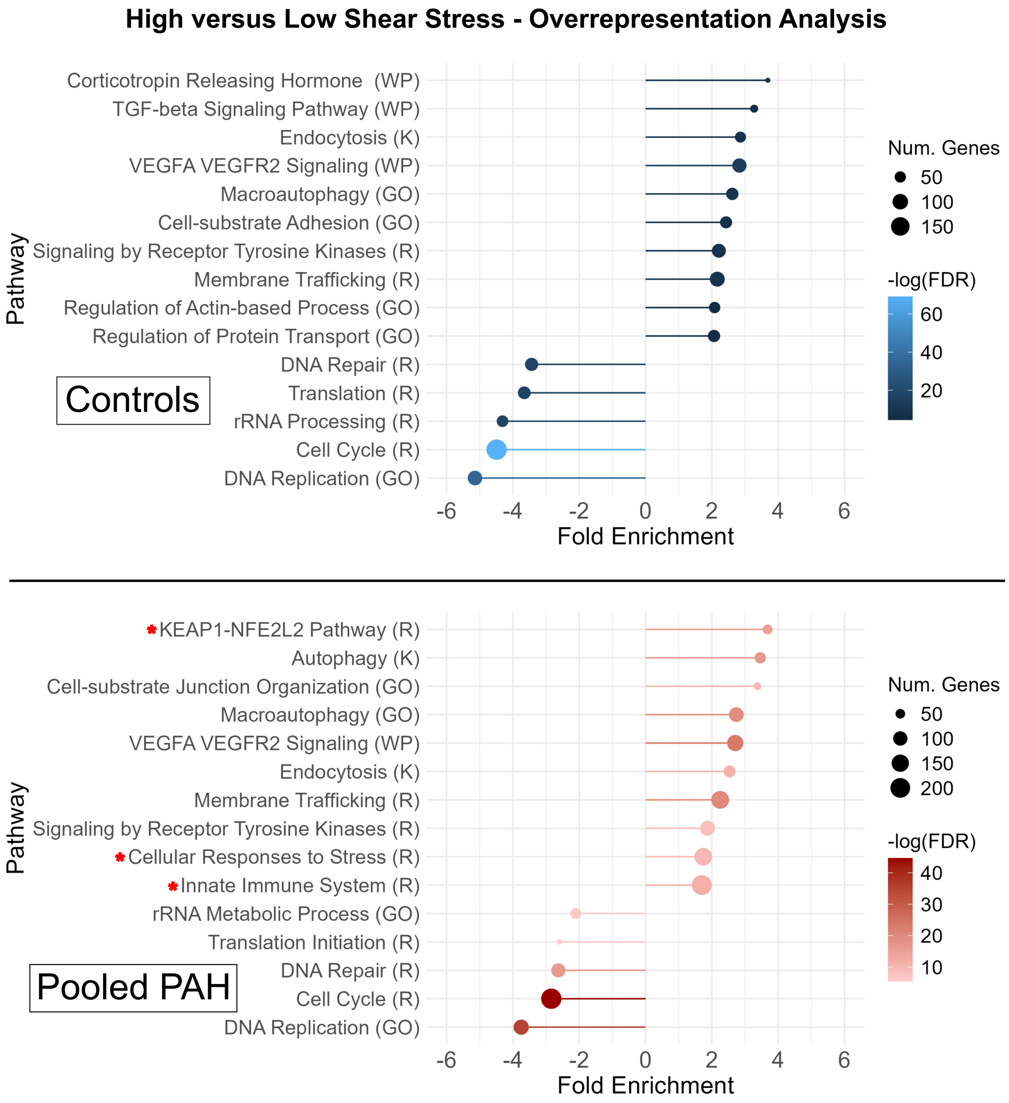


**Figure S3: Overrepresentation analysis of PAEC signaling in response to high versus low shear:** Overrepresentation analysis was performed on control PAECs (n = 3; top, blue) and pooled PAH PAECs from donors with idiopathic or congenital heart disease-PAH (n = 6; bottom, red) under high versus low shear, with selected pathways displayed here. Positive fold enrichment signifies pathways enriched under high shear – negative fold enrichment signifies those enriched under low shear. PAH cells displayed increased enrichment for pathways related to cellular stress or immune signaling (red asterisk). Significance was determined by an FDR q-value <0.05. Pathways are annotated with gene sets from Hallmark (H), Kyoto Encyclopedia of Genes and Genomes (KEGG), Reactome (R), WikiPathways (WP), and Gene Ontology Biological Processes (GO), with pathway names abbreviated where appropriate.

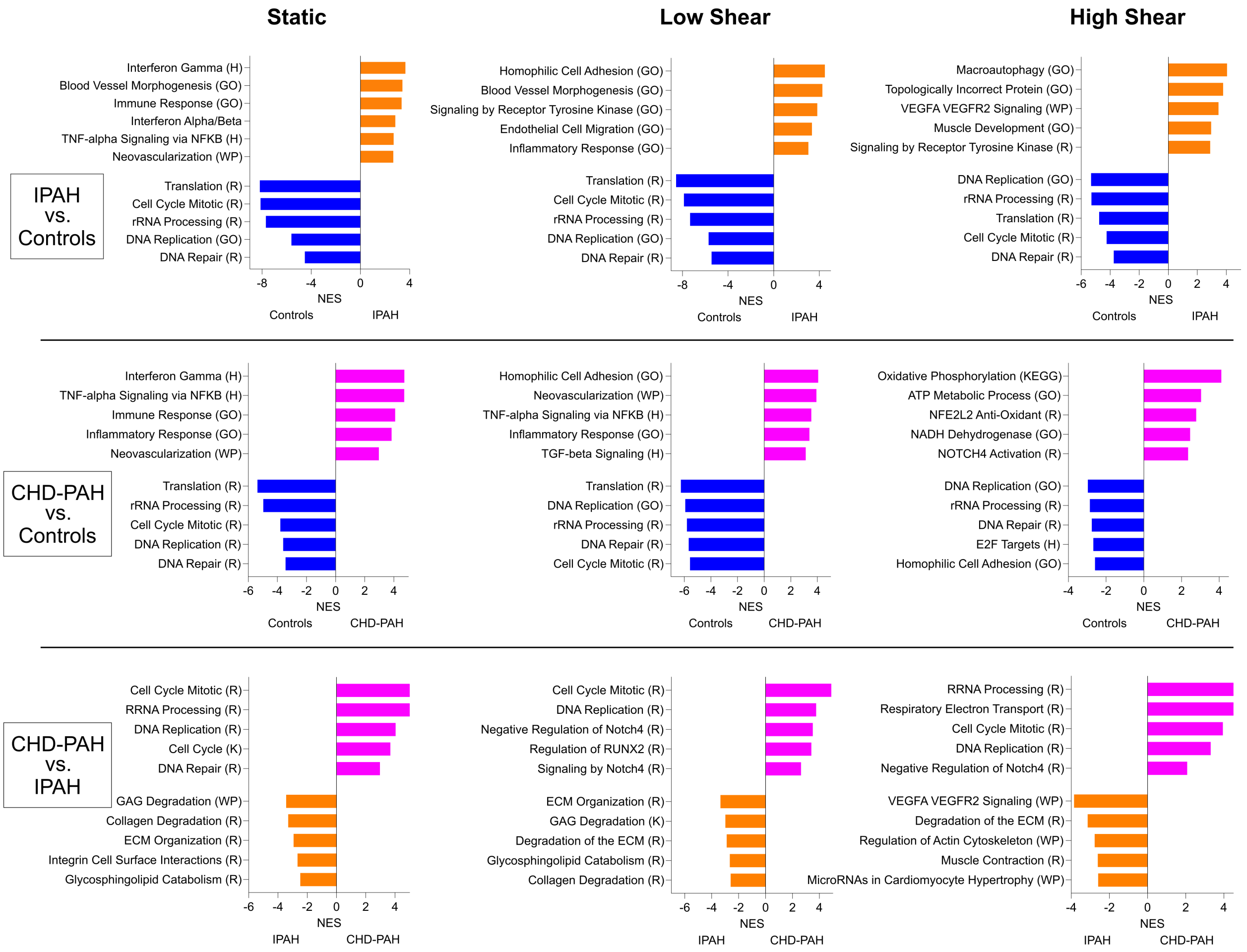


**Figure S4: Gene set enrichment analysis (GSEA) of PAH subtypes under shear**. Selected significantly enriched pathways comparing IPAH vs Controls, CHD-PAH vs Controls, and CHD vs IPAH across three shear conditions. Significance was determined by an FDR q-value <0.05 and the degree of enrichment represented by the normalized enrichment score (NES). Pathways are annotated with gene sets from Hallmark (H), Kyoto Encyclopedia of Genes and Genomes (KEGG), Reactome (R), WikiPathways (WP), and Gene Ontology Biological Processes (GO), with pathway names abbreviated where appropriate.

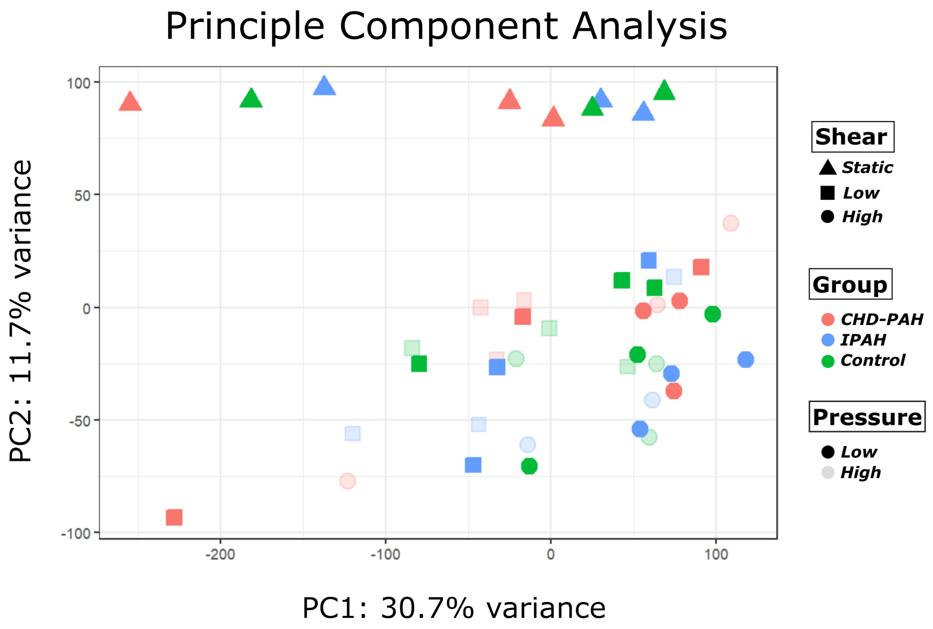


**Figure S5: Principal component analysis of pressure analysis**: Data showing transcriptional variance for a subset of donors studied under high pressure vs low pressure conditions across a range of shear stress (n = 3 for each group with one sample excluded as an outlier in the IPAH high shear, high pressure group). Shape denotes shear state, color defines patient group, and transparency indicates pressure condition (solid = low, transparent = high).

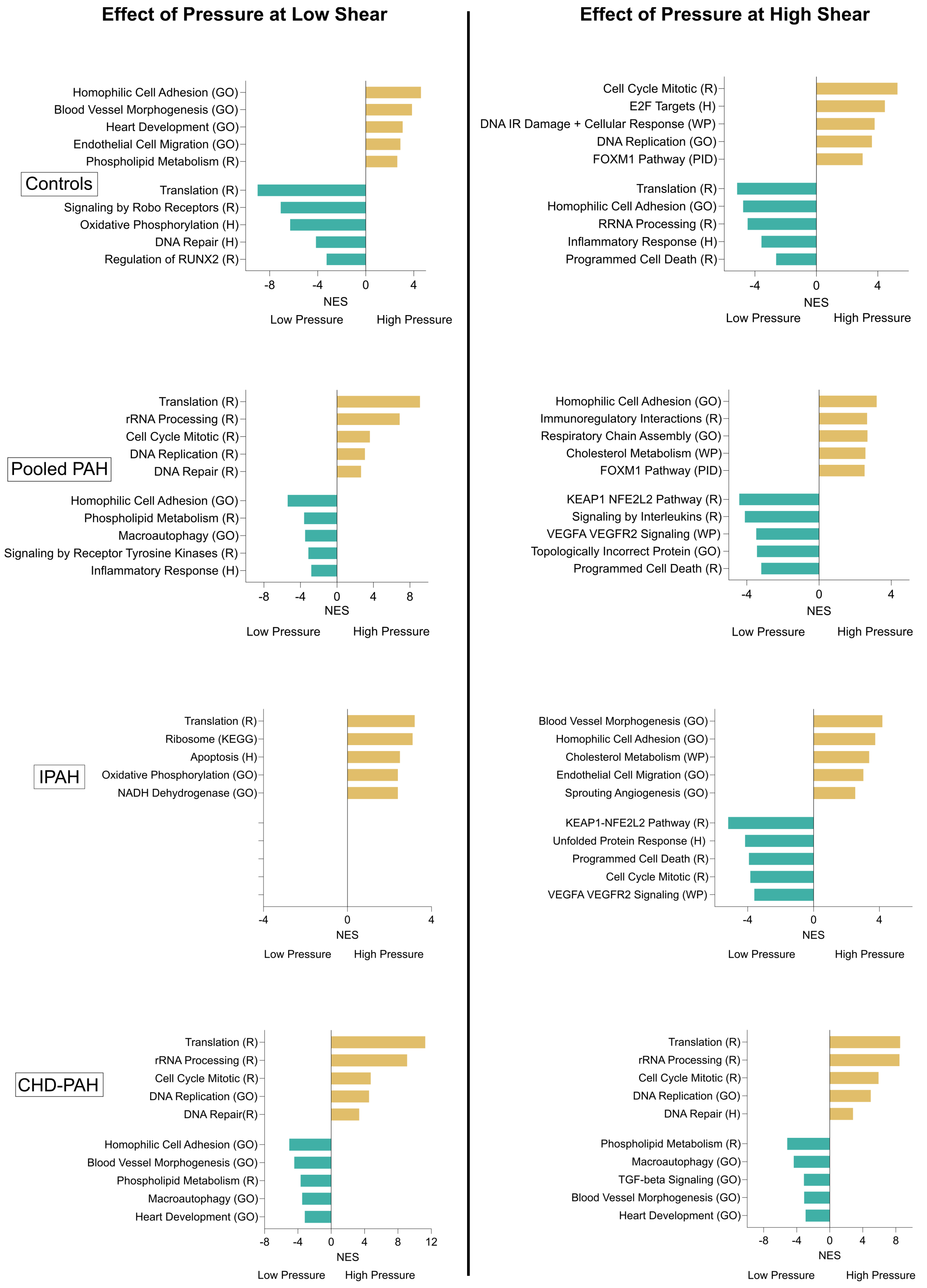


**Figure S6: Differential transcription under high vs low shear across subject groups and shear states.** Gene set enrichment analysis was used to identify significantly enriched pathways under high pressure versus low pressure conditions across subject groups and at low (left) and high (right) shear. Significance was determined by an FDR q-value <0.05 and the degree of enrichment represented by the normalized enrichment score (NES). Pathways were annotated with gene sets from Hallmark (H), Kyoto Encyclopedia of Genes and Genomes (KEGG), Reactome (R), WikiPathways (WP), and Gene Ontology Biological Processes (GO), with pathway names abbreviated where appropriate.

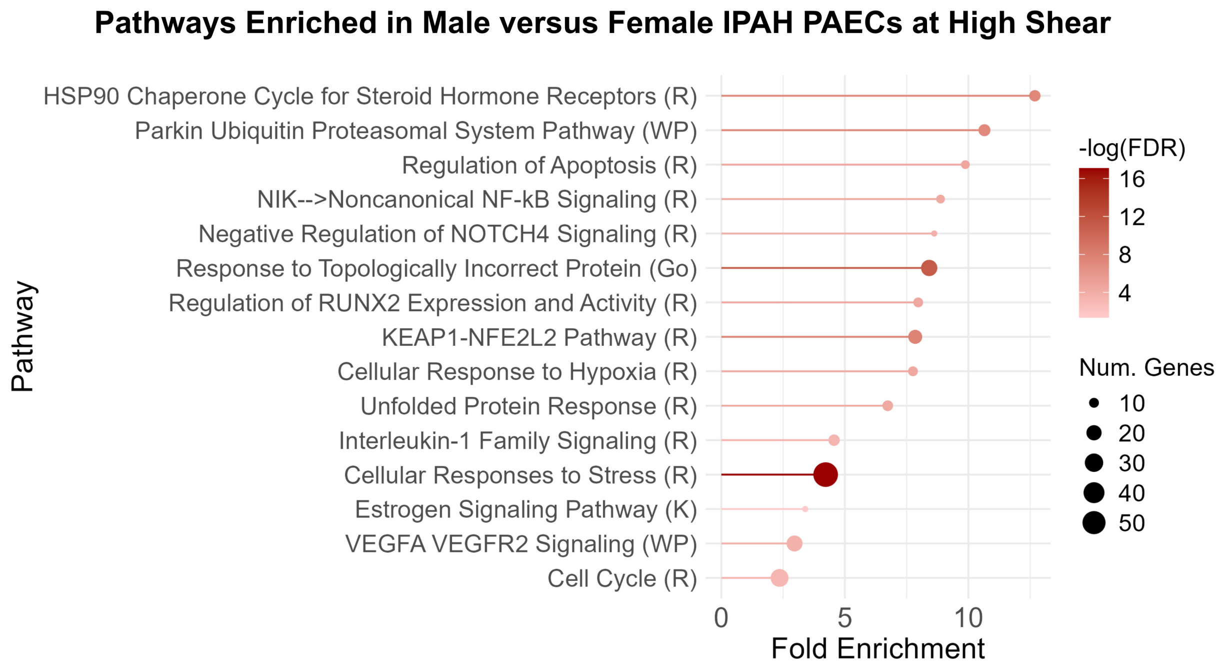


**Figure S7: Overrepresentation analysis of male vs female IPAH PAECs under high shear:** Overrepresentation analysis was performed on male vs female PAECs from donors with IPAH (n = 3 for each group) under high shear stress, with selected pathways displayed here. Male PAECs displayed increased enrichment of pathways involved in cellular stress responses, immune signaling, proliferation, and protein degradation, as well as differences in vascular cytokine and sex-steroid signaling. Significance was determined by an FDR q-value <0.05. Pathways were annotated with gene sets from Hallmark (H), Kyoto Encyclopedia of Genes and Genomes (KEGG), Reactome (R), WikiPathways (WP), and Gene Ontology Biological Processes (GO), with pathway names abbreviated where appropriate.
