## Supplementary Methods for "Pressure Points: Endothelial Responses to Shear Stress and Pressure in Health and Pulmonary Arterial Hypertension"

Design and fabrication of the resistor-coupled microfluidic: Culture channels (500 µm wide, 112 µm in height, and 20 mm long) were fabricated by casting PDMS against a silicon mold. Silicon molds were fabricated in a cleanroom using standard lithography techniques. Channels were designed using LayoutEditor (juspertor GmbH) and transferred to a chrome mask via exposure on a DWL 66+ (Heidelberg GmbH) followed by etching. Silicon wafers were then coated with AZ9260 negative photoresist (Microchem) to a thickness of 6 µm, aligned to the above chrome mask using a contact photolithography aligner (ABM), exposed, developed using AZ400K (Microchem), and subjected to deep reactive ion etching (SPTS Rapier DRIE) followed by resist removal (EKC, Dupont). Dimensions were confirmed with a non-contact profilometer (Keyence VK-X150K) as well as a contact profilometer (KLA Tencor P15). Wafers were then silanized with trichloro (3,3,3- trifluoropropyl) silane (Sigma). To form PDMS channels, a 10:1 mixture of base and curing agent (Sylgard 184, Dow) was thoroughly mixed, degassed, poured over the patterns, and cured at 65 °C for 3 hours or more. Channels were excised using a scalpel and inlet and outlet holes created with a 2 mm biopsy punch (Integra Miltex). Large glass slides (75.5mm x 50.4mm x 1mm, Ted Pella) were cut using a glass cutter to make glass platforms to serve as a base for PDMS channels. To facilitate cellular adhesion, glass slides were cleaned in 1M hydrochloric acid (50-60°C) for 8-12 hours followed by washes with distilled water and 100% ethanol and blot drying with bibulous filter paper (Whatman). Patterned PDMS channels were then bonded to these treated glass platforms using air plasma (75 W, Plasma Etch), edges further secured to the glass with a thin brushed-on layer of 10:1 PDMS mixture as above, followed by annealing in a 65 °C oven for at least 1 hour. Microfluidic resistors were created via needle-subtraction technique – casting a 100 µm-diameter wire in PDMS, with the wire then removed, inlets/outlets punched at the desired length, and the entry/exit of the wire past the biopsy punched areas carefully sealed with PDMS. For perfusion experiments, 3.5% dextran-containing culture media was used, to simulate the viscosity of blood (~3.5 cP).

Hemodynamic conditions: The total pressure range across culture channels was calculated as the sum of the pressure generated by the microfluidic resistor and the intrinsic resistance of the culture channel itself. The pressure drop across resistors (circular cross-section) was calculated using the Hagen–Poiseuille equation $\Delta P = \frac{8\mu QL}{\pi R^4}$ where $\Delta P$ reflects the pressure drop, $\mu$ dynamic viscosity, $Q$ the volumetric flowrate, $L$ the resistor length and $R$ the radius. The pressure drop across the culture channel (rectangular cross-section) was calculated using $\Delta P = \frac{16\pi^2 \mu QJL}{(hw)^2}$ where $J$ refers to the polar moment of inertia for a rectangular cross-section: $J = \frac{hw(h^2+w^2)}{12}$ as derived from prior work.[1] Shear stress was calculated as $\tau = \frac{6Q\mu}{h^2 w}$ in dyne/cm$^2$. Flow rates and resistor lengths were calculated to provide the desired hemodynamic parameters (**Table 1, main text**). Pressure varied slightly along the culture channel, as tabulated, due to the resistance imposed by the culture channel itself. Verification that desired pressure was achieved was conducted by perfusion using a microfluidic pump controller and both flow and pressure sensors, as previously described.[2]

Preparation and perfusion of culture channels: To culture cells in PDMS chambers prepared as above, channels were sterilized via autoclave. To enhance cell adhesion they were then coated with 0.1% polydopamine[3] by perfusing an alkalinized solution of dopamine through the channels. Dopamine hydrochloride powder (Sigma-Aldrich) was diluted to 0.1% by mass into 10 mM Tris-HCL (pH 8.5), rapidly syringe filtered, and immediately perfused into channels which were then placed into a dark chamber and left at room temperature for one hour. From this point on, channels were not allowed to dry and kept continuously in aqueous solution, with great care taken not to introduce any bubbles into microfluidic channels. Unbound polydopamine was removed by washing with cell-grade water. Channels were then perfused with a solution of 5 µg/mL fibronectin in PBS and incubated at 37°C for at least one hour before washing with 37°C culture media to prepare for cell seeding. PAECs were collected, resuspended at a concentration of 4 $*10^6$ cells/ml, and 10 µL of this solution injected into culture channels. Cells were allowed to adhere for 1 hour at 37°C followed by rinsing with culture media to remove unbound cells. Importantly, cells in the inlet and outlet reservoir were physically removed by scraping with an aspirating pipette tip hooked up to suction, leaving cells only in the channels themselves where developed flow profiles are expected. Cells were allowed to attach/proliferate overnight in a 37°C incubator using intermittent slow flow obtained by inserting media-filled pipette tips into the inlet and outlet and leaving channels on a custom rocking platform set to alternate between -8 and +8 degrees of tilt every hour.

Morphologic analysis: Following 24 hours of flow exposure, culture chambers were disconnected from the syringe pump and brightfield imaging was rapidly obtained for morphologic analysis prior to immediately moving to RNA isolation (below). Images were obtained using phase-contrasted brightfield microscopy with a 20x objective, and imported into the Fiji distribution of ImageJ software (U.S. National Institute of Health, Bethesda, MD).[4,5] A defined 500 µm x 1000 µm rectangle was selected for analysis, away from the inlet or outlet. Cell borders were manually traced and measurements taken in Fiji, including cell shape indices and the angle of orientation relative to the direction of flow. Cells were considered “elongated” if their circularity index was ≤ 0.7, and elongated cells oriented within ± 30° of the direction of flow were considered flow-aligned as well, as previously described.[6] The percentage of cells that were elongated and flow-aligned was compared across conditions via ANOVA using Graphpad Prism software v9.5.1 (GraphPad Software, San Diego, CA, USA), with Šidák post-hoc test performed for multiple comparisons.

Immunofluorescent staining and imaging: To verify endothelial cell markers, as well as to better visualize alignment in chambers, staining was performed on both static cultures and culture chambers that were not used for RNA collection. Cells or chambers were fixed in 4% Paraformaldehyde solution for 15 minutes, followed by permeabilization and blocking for 1 hour in 0.1% triton in 2% BSA, staining with desired antibodies for 1 hour at room temperature, washing with PBS followed by optional staining with a secondary antibody for 1 hour at room temperature and additional washing with PBS (for non-conjugated antibodies), and imaging on a Nikon A1R confocal microscope. Antibodies used were: Alexa Fluor 488-conjugated rabbit anti-

VWF (1:100 Abcam ab195028), rabbit anti-VE-cadherin (1:50, Abcam ab33168), Alexa Fluor 568-conjugated Phalloidin (1:100, Thermo Fisher A12380), goat anti-rabbit Alexa Fluor 647 (1:100, Thermo Fisher A21244), and Hoechst (1:250, Thermo Fisher) for nuclear staining.

RNA sequencing analysis: The following pipeline was used to analyze RNA sequencing data after raw read alignment and gene count generation.

*1. Filtering and batch correction*: Raw gene counts were filtered to exclude genes with <0.5 counts per million reads. To correct for batch effects due to running sequencing in two batches, ComBat-seq (sva package, version 3.48.0) was run in R (version 4.3.1).[7] One IPAH donor in the high pressure, high shear condition was removed a-priori as a clear technical outlier.

*2. Principal-component analysis (PCA) and heatmap*: To visualize data under low-pressure conditions, PCA was performed in R using the prcomp function with scaling enabled. Preprocessing was done by filtering out non-count columns and eliminating genes with zero variance across all samples. For further global data visualization under low pressure, a heatmap was prepared using the IDEP interface (version 1.1, via https://bioinformatics.sdstate.edu/idep/).[8] To generate a heatmap of gene expression by sample, the 1000 most variable genes were included, using correlation for distance, average linkage, a Z-score cutoff of 3, with genes centered and normalized.

*3. Differential gene expression*: Differentially expressed genes were identified with DESeq2 package (version 1.40.2) in R. When the same cell lines were compared across different hemodynamic conditions, paired analyses were performed. Adjusted p-value <0.05 was considered significant.

*4. Gene set enrichment analysis*: Pathway enrichment in response to hemodynamic stimuli was assessed using Gene Set Enrichment Analysis (GSEA) software version 4.3.2 (Broad Institute, USA) in pre-ranked mode. All genes were ranked by Wald statistic ($\log_2$ fold change divided by its standard error) derived from DESeq2 (version 1.40.2) in R. The following gene set collections were tested: Hallmark (h.all.v2023.2.Hs.symbols.gmt), Biocarta (c2.cp.biocarta.v2023.2.Hs.symbols.gmt), KEGG (c2.cp.kegg.v2023.2.Hs.symbols.gmt), Pathway Interaction Database (PID, c2.cp.pid.v2023.2.Hs.symbols.gmt), Reactome (c2.cp.reactome.v2023.2.Hs.symbols.gmt), WikiPathways (WP, c2.all.v2023.2.Hs.symbols.gmt), and Gene Ontology Biological Processes (GO, c5.go.bp.v2023.2.Hs.symbols.gmt). Enrichment was evaluated using the classic scoring scheme with 1000 gene set permutations. Only gene sets containing between 5 and 500 genes were included. Pathways were considered significantly enriched at FDR q-value < 0.05. Enrichment strength was quantified as a normalized enrichment score (NES), as computed by GSEA.

*5. Over-representation analysis*: Genes determined to be differentially expressed (adjusted p-value <0.05) between two hemodynamic conditions underwent functional assessment via over-representation analysis (ORA). Pathways were annotated with GO Biological Processes, KEGG, Reactome, and WP with significance determined by FDR <0.05. Degree of pathway overrepresentation was quantified by the ratio of measured genes divided by expected genes in each respective pathway. ORA was performed with the WEB-based GEne SeT AnaLysis Toolkit

(WebGestalt [v2019], via https://www.webgestalt.org/) using Benjamini-Hochberg correction and a minimum number of 5 analytes per category.[9]

*6. Enrichment map*: Cytoscape (version 3.10.2) EnrichmentMap App[10] was used to illustrate enriched gene sets, identified by GSEA, as a network. Visualization was organized so that nodes represent gene sets, with node size proportional to the absolute value of NES, and interconnecting lines or edges representing genes shared between the gene sets with thickness proportional to the number of overlapping genes. Node and edge cutoffs of q-value 0.1 and 0.5 were selected, respectively, for illustration purposes.

*7. Short Time Series Expression Miner (STEM) Analysis*: STEM (version 1.3.13) evaluates how gene expression changes with respect to an increasing independent variable such as time.[11] In this case, the gene expression of endothelial cells was evaluated across escalating conditions of shear (static, low, and high shear) as expression curves transformed so that the static condition value was set equal to zero. Gene expression curves were matched to predefined model expression profile curves by correlation coefficients using a significance cutoff of FDR < 0.05. We were interested in global endothelial responses to shear under the low-pressure condition, and thus all donors were included for STEM analysis. After filtering and batch correction (above), counts for each gene at each shear condition were averaged across all donors and $\log_2$ normalized. The STEM clustering method was used. The maximum number of model profiles was 50 and the maximum unit change in model profiles between shear conditions was 2. The maximum number of candidate model profiles was 1,000,000 and number of permutations per gene was 50. For matching gene expression curves to predefined expression curves, a minimum correlation coefficient of 0.7 was required. Genes matched to each statistically significant expression profile were carried forward with an overrepresentation analysis and functionally annotated based on GO Biological Process, KEGG, Reactome, and WP using WebGestalt [2019 version] as above. An FDR cutoff of <0.05 was used for pathway significance.

**Supplementary References**:


1. Bahrami M, Yovanovich MM, Culham JR. Pressure drop of fully-developed, laminar flow in microchannel of arbitrary cross-section. *Journal of Fluids Engineering, Transactions of the ASME*. 2006;128(5):1036–1044.
2. Mandrycky C, Ishida T, Rayner SG, Heck AM, Hadland B, Zheng Y. Under pressure: integrated endothelial cell response to hydrostatic and shear stresses. *bioRxiv : the preprint server for biology*. 2024.
3. Ding YH, Floren M, Tan W. Mussel-inspired polydopamine for bio-surface functionalization. *Biosurface and biotribology*. 2016;2(4):121–136.
4. Schneider CA, Rasband WS, Eliceiri KW. NIH Image to ImageJ: 25 years of image analysis. *Nature methods*. 2012;9(7):671–5.
5. Schindelin J, Arganda-Carreras I, Frise E, Kaynig V, Longair M, Pietzsch T, Preibisch S, Rueden C, Saalfeld S, Schmid B, Tinevez J-Y, White DJ, Hartenstein V, Eliceiri K, Tomancak P, et al. Fiji: an open-source platform for biological-image analysis. *Nature*

*methods*. 2012;9(7):676–82.

6. Wang C, Baker BM, Chen CS, Schwartz MA. Endothelial cell sensing of flow direction. *Arteriosclerosis, thrombosis, and vascular biology*. 2013;33(9):2130–6.

7. Zhang Y, Parmigiani G, Johnson WE. ComBat-seq: batch effect adjustment for RNA-seq count data. *NAR genomics and bioinformatics*. 2020;2(3):lqaa078.

8. Ge SX, Son EW, Yao R. iDEP: an integrated web application for differential expression and pathway analysis of RNA-Seq data. *BMC bioinformatics*. 2018;19(1):534.

9. Liao Y, Wang J, Jaehnig EJ, Shi Z, Zhang B. WebGestalt 2019: gene set analysis toolkit with revamped UIs and APIs. *Nucleic acids research*. 2019;47(W1):W199–W205.

10. Shannon P, Markiel A, Ozier O, Baliga NS, Wang JT, Ramage D, Amin N, Schwikowski B, Ideker T. Cytoscape: a software environment for integrated models of biomolecular interaction networks. *Genome research*. 2003;13(11):2498–504.

11. Ernst J, Bar-Joseph Z. STEM: a tool for the analysis of short time series gene expression data. *BMC bioinformatics*. 2006;7:191.
